# Living Bacterial Reservoir Computers for Information Processing and Sensing

**DOI:** 10.1101/2024.09.12.612674

**Authors:** Paul Ahavi, Thi-Ngoc-An Hoang, Philippe Meyer, Sylvie Berthier, Federica Fiorini, Florence Castelli, Olivier Epaulard, Audrey Le Gouellec, Jean-Loup Faulon

## Abstract

We introduce a systems-level approach to sensing and computing in which *Escherichia coli* acts as a living reservoir computer, performing complex information processing through its native growth responses without requiring genetic modification or specialized instrumentation.

We validate this framework by accurately classifying early-stage COVID-19 plasma samples (mild *vs*. severe) using only bacterial growth data, highlighting a diagnostic potential without infrastructure-dependent methods. By controlling nutrient media compositions, we also demonstrate that *E. coli* growth encodes nonlinear transformations that outperform linear regression, support vector machines, and multilayer perceptrons across diverse regression and classification tasks. Using simulations across genome-scale metabolic models from multiple bacterial species, we establish a strong link between phenotypic diversity and computational capacity, showing that learning capacities scale with the diversity of metabolic phenotypes.

These findings position biological reservoir computing as a robust, scalable, and low-cost platform for intelligent biosensing, diagnostics, and hybrid bio-digital computation, while providing new mechanistic insights into the computational capabilities of living systems.

## Introduction

Sensing and computing with living organisms is one of the main endeavors of synthetic biology where many electronic-like circuits have been engineered in a bottom-up fashion^1,2^. Engineering and tuning such circuits is challenging and several factors such as metabolic burden^3^, orthogonality issues^4^ and low noise tolerance^5^ strongly restrain their complexity.

Synthetic biology devices have largely been inspired by behaviors observed in natural systems. Indeed, information processing systems analogous to logic gates^6^, switches^7^, and perceptrons^8^ have all been discovered in various organisms. This raises the question: *instead of engineering bottom-up devices, could we develop a top-down approach where natural organisms are repurposed to perform complex computational tasks*?

The study of physical systems’ ability to solve computational tasks falls within the field of Artificial Intelligence known as reservoir computing (RC)^9^. RC systems come in two broad forms: conventional and physical RC. Conventional RC systems are *in silico* tools, whereas physical RC uses real physical systems as reservoirs.

Physical RC begins by presenting the task input, typically encoded as a physical stimulus, to the physical reservoir, which maps it to a high-dimensional, time-evolving state. These states are then passed to a readout, usually a simple linear classifier or regressor, that produces the final output. Physical RC has many applications, including the seminal liquid state machine for pattern recognition in a bucket of water^10^, chemical RC developments for classification tasks and time-series forecasting^11^, and RC systems using rat cortical neurons cultured on micropatterned substrates to solve classification tasks^12^.

As illustrated in Fig. 1, the aim of this study is to use *E. coli* (K-12 MG1655) as a sensing and reservoir computing platform. We first present a practical application by predicting COVID-19 disease severity (mild vs. severe) from plasma samples, and then show how our *E. coli* reservoir can be leveraged as a physical RC system to solve several benchmark machine-learning (ML) classification and regression tasks. We conclude by establishing a relationship between strain phenotypic diversity and problem-solving performance.

**Fig. 1.**
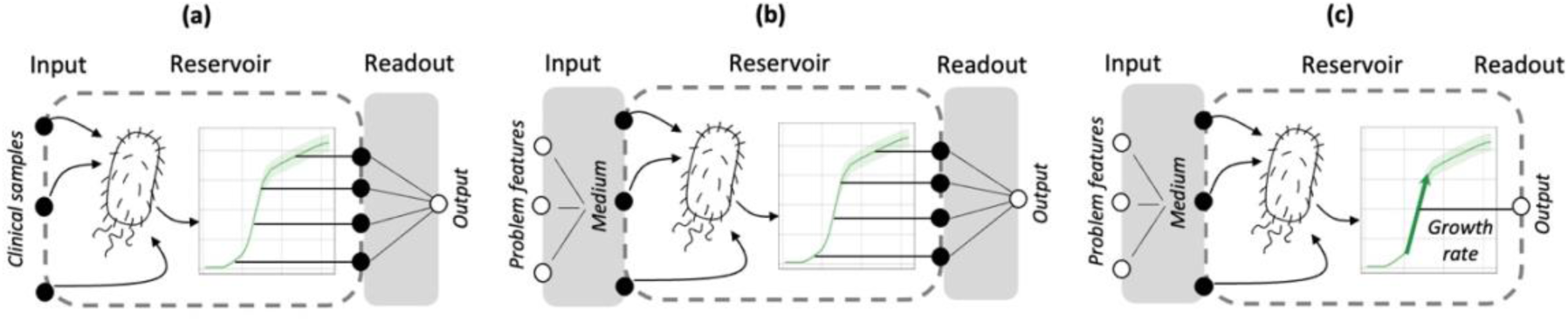
Bacterial reservoir computing (RC) frameworks. The Figure, inspired by Tanaka et al.^9^, was adapted for bacteria. Gray regions indicate the trainable components of each framework. **(a)** Bacterial RC for sensing. E. coli is grown on plasma samples (COVID-19 mild or severe), and growth curves are recorded. The readout is an external classifier that takes optical density (OD) measurements at multiple time points to predict disease class (mild or severe). **(b)** Bacterial RC for solving machine-learning (ML) problems using growth curves. ML input features are transformed into a growth medium. E. coli is grown in this medium and the growth curve is recorded. The ML task output is computed by an external trainable linear classifier or regressor. **(c)** Bacterial RC for solving ML problems using growth rates. As in panel (b), ML input features are transformed into a growth medium. E. coli is grown in this medium and the growth rate is measured. No readout is trained: the normalized growth rate is used directly to predict the normalized ML target.

## Results

### Classifying plasma samples with an *E. coli* reservoir

Classifying COVID-19 patients to predict whether they will develop a mild or severe form of the illness has been a significant research focus since the onset of the pandemic^13^. Most methods performing classification are making use of proteomic and metabolomic profiling conducted on biological samples^13,14,15^ collected from patients upon their admission to the hospital. After having been trained on these omics data, ML (*cf*. Abbreviations in Materials and Methods **M1**) techniques are used to predict the disease outcome (e.g., mild or severe according to whether ICU admission was required^14,16^).

In line with this approach, we utilized a cohort of COVID-19 patients from the CHUGA University Hospital (Materials and Methods **M2**) and performed untargeted metabolomics analyses (via high resolution mass spectrometry) on plasma samples collected on 81 patients (50 mild, 31 severe, *cf.* Supplementary Methods **MS1**). These samples were collected before they developed either mild or severe forms of the disease. We observe that disease outcomes can be predicted from these metabolomics data, with cross-validation accuracies greater than 0.9 (Supplementary Methods **MS2** and Figure **S1**). These accuracies were obtained by selecting MS signals maximizing classification accuracies (Supplementary Methods **MS3**). Because our goal is to develop a physical reservoir for classifying the samples, we also generated predictions using only signals corresponding to metabolites expected to cross the *E. coli* membrane and contribute to its growth. The results shown in Supplementary Figure **S1** show accuracies above 0.8, indicating that metabolites crossing the *E. coli* membrane are sufficient to distinguish mild from severe cases. This suggests that these metabolites could differentially influence *E. coli* growth motivating further development of a physical reservoir.

Seeking to construct an *E. coli* reservoir, we added plasma samples from COVID-negative patients, obtained from the French Blood Establishment (Etablissement français du sang, EFS) to the mild and severe samples extracted from the CHUGA COVID-19 cohort (Materials and Methods **M2**). Since *E. coli* cannot grow in the presence of antibiotics, any samples containing antibiotics were excluded. The final dataset consisted of 27 negative samples, 37 mild cases, and 36 severe cases. All these samples had to be pretreated in order to optimize bacterial growth on plasma and therefore classification potential across sample categories (Materials and Methods **M3**).

By growing *E. coli* on negative, mild, and severe plasma samples, we first found out that an *E. coli* RC system relying solely on growth rate as input to the post-readout classifier was unable to solve the mild *vs*. severe classification task. We also observed we could not classify mild *vs*. severe when taking lag time and maximum optical density taken at 600 nm (ODMAX) as readout input in addition to growth rates (Supplementary Table **S1**). To further increase the number of features available to the readout classifier, we utilized optical density (OD) measurements taken at all time points along the growth curve as in **Fig. 1a**.

Using the framework shown in **Fig. 1a**, and 20-fold cross-validation for the readout classification, our *E. coli* RC system achieved a classification accuracy of 0.86 for distinguishing mild from severe samples (**Fig. 2**). This accuracy is similar to the one obtained using MS data for metabolite crossing the *E. coli* membrane (Supplementary Figure **S1**). Furthermore, the reservoir demonstrated high accuracy (0.95) in distinguishing negative samples from positive ones. As with MS data, these results were obtained by selecting sets of time points that maximized classification accuracies (Supplementary Methods **MS3**). Results for other classifiers (SVM and MLP) and other folds (5 and LOO) are presented in Supplementary Table **S2**.

**Fig. 2.**
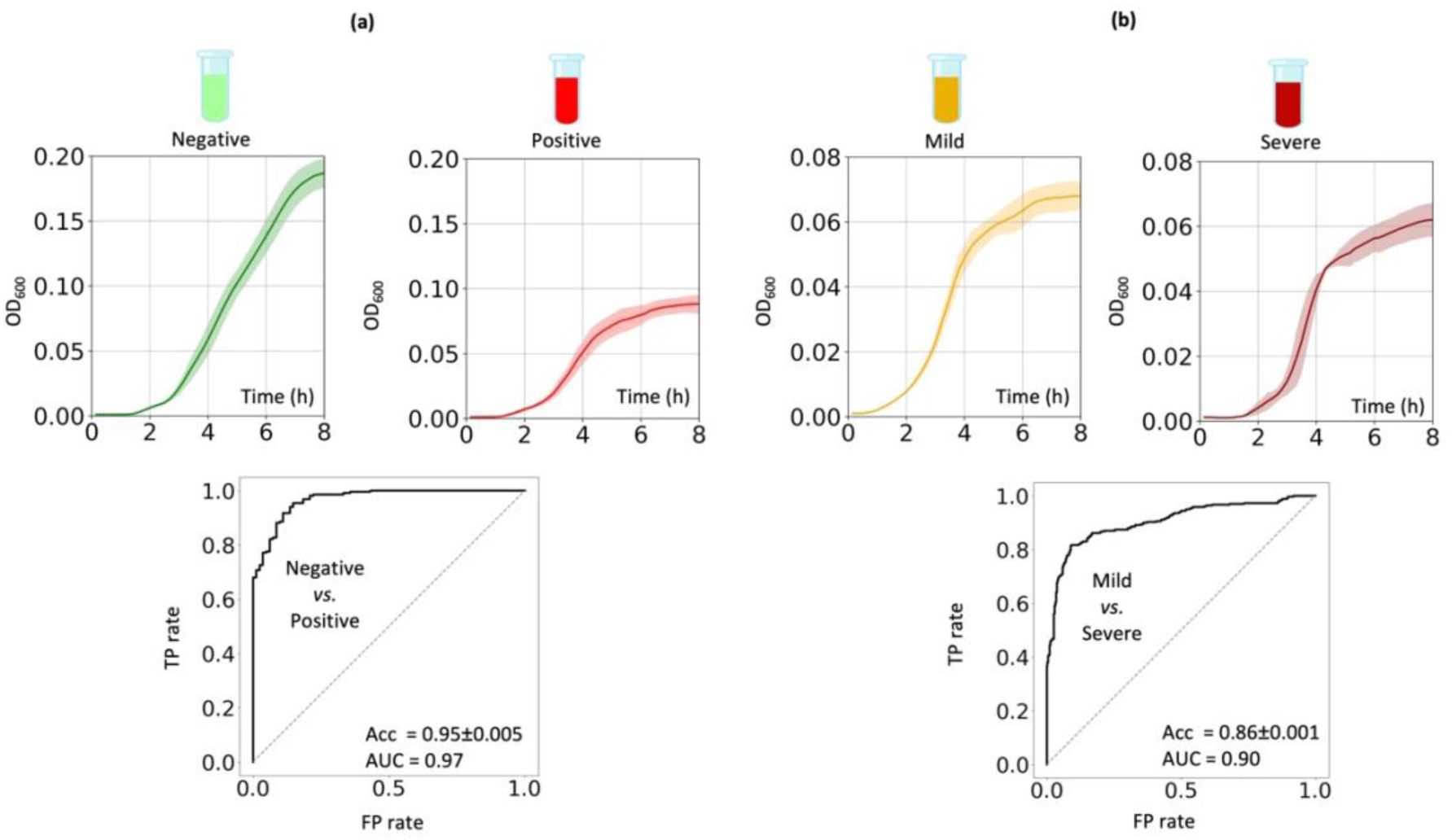
Classifying COVID-19 plasma samples with E. coli RC. The RC framework is shown in Fig. 1a. **(a)** Classifying COVID-19 negative vs. positive plasma samples. The input is a negative or positive plasma sample. The reservoir is MG1655 on which growth curves (2 technical replicates) are acquired using a plate reader recording OD_600_ at 48 time points taken from 0 to 8h (cf. Materials and Methods **M3**). The readout is a XGB classifier that takes as input OD_600_ values for selected time points and prints out an accuracy for 20-fold cross validation repeated three times. Growth curves are shown for sample N1 from Supplementary Figure **S2** and sample M10 from Supplementary Figure **S3**. **(b)** Classifying COVID-19 mild vs. severe plasma. The set up is identical to panel a. Growth curves are shown for sample M9 from Supplementary Figure **S3** and sample S8 from Supplementary Figure **S4**. Growth curves for all samples are plotted in Supplementary Figures **S2-4** and all OD_600_ values can be found in the supplementary file ‘Reservoir_Covid’.

For the mild vs. severe classification task, the selected time points were mostly within the first 6 hours. In contrast, for the negative vs. positive classification, the selected time points extend up to 8 hours. Generally, *E. coli* biomass at 8 hours is lower in positive samples compared to negative ones, and we observe that growth is generally delayed in the first 4 hours when *E. coli* is grown on severe samples compared to mild ones. These finding are substantiated by an analysis we performed using Generalized Additive Mixed Models (GAMMs^17^, *cf.* Materials and Methods **M4** and Supplementary Table **S3**). GAMMs confirmed statistically significant differences in growth dynamics between groups. Negative and positive samples differed in both mean OD and temporal trajectory, whereas mild and severe samples differed primarily in the shape of the OD time course (with delayed early growth in severe cases), despite similar mean OD levels.

### Bacterial reservoir computing framework to solve machine-learning problems

Because we observed *E. coli* can act as a reservoir to classify plasma sample, we investigated whether this capability could be leveraged in a reservoir computing framework to solve ML problems. In typical ML regression and classification tasks, training sets are composed of features (input) and labels (output), with the goal of predicting the labels from the features. As shown in **Fig. 1b–c**, solving an ML task with our *E. coli* reservoir starts by converting each ML feature vector into an experimental condition, *i.e.*, choosing a growth medium on which our *E. coli* reservoir is grown.

The challenge is to find the most appropriate mapping between features and nutrients. In the reservoir computing (RC) literature, there is no generic procedure for feeding problem features into a reservoir; instead, different methods are used depending on the physical system (*e.g*., using water waves to mimic sound waves for classifying voice recordings in liquid state machines^10^). In our case, one possible strategy could be an arbitrary one-to-one mapping of features to nutrients. However, this approach may not yield the best performance. Alternatively, we can optimize the nutrient inputs by training a features-to-medium mapper that selects nutrients (**Fig. 3a** and Materials and Methods **M5)**. Although many RC implementations use a fixed input mapping, training a reservoir input is possible and has been explored previously^18,19^.

**Fig. 3.**
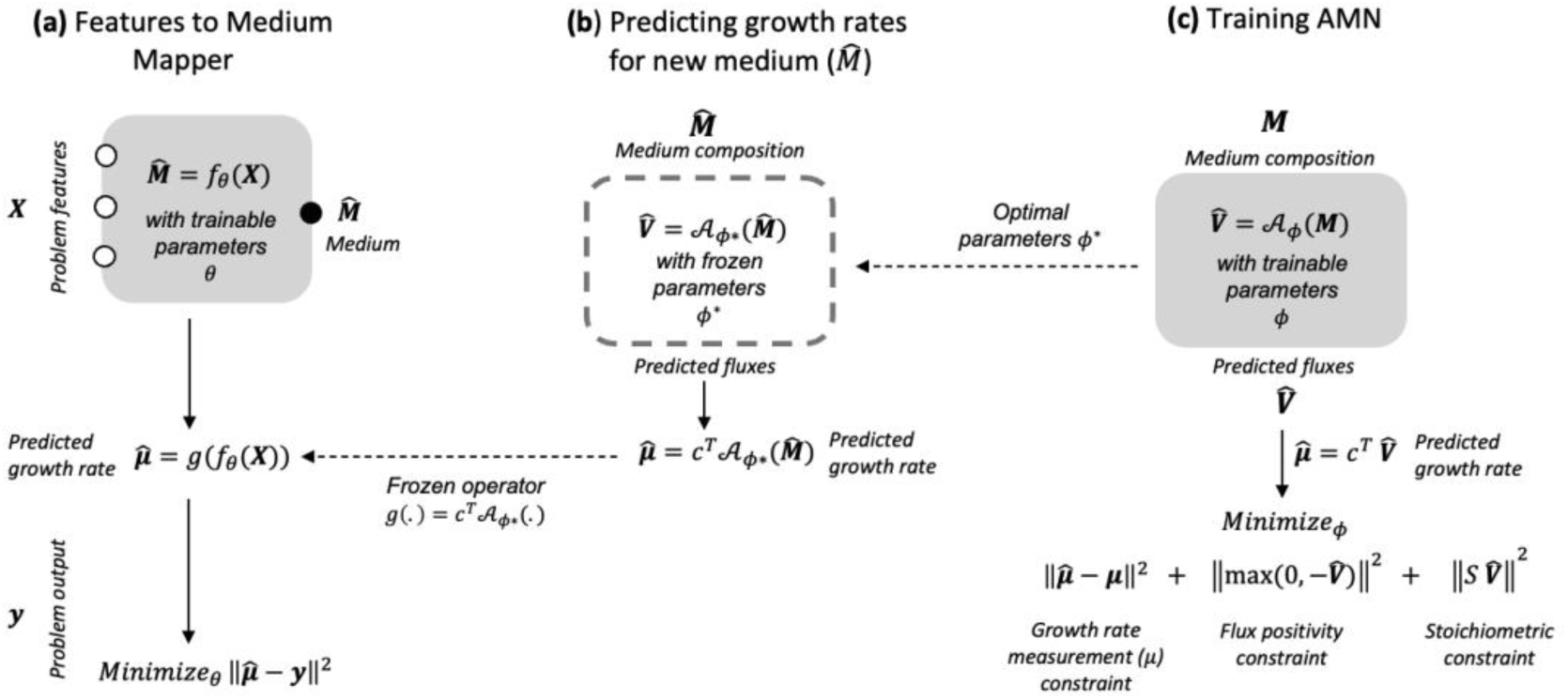
Features-to-medium workflow. **(a)** The features-to-medium mapper is a trainable function, *f*_θ_, predicting a medium 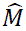 from a feature vector X. The mapper *f*_θ_ is an MLP followed by a softmax layer that selects one medium among 280 experimentally prepared media (details in Materials and Methods **M5**). It uses a frozen operator *g*(.), whose role is to predict the growth rate 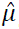 for any candidate medium 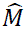. Because both the predicted growth rate 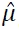 and the problem output y are normalized, *f*_θ_ is trained by minimizing their difference. **(b)** Fluxes 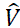 are predicted using the trained AMN 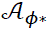, whose parameters ϕ^∗^ are frozen and used across all ML tasks. The operator *g*(.) is obtained by extracting the predicted growth rate 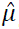 from the predicted reaction flux vector 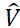 using the biomass projection 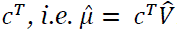. **(c)** The AMN 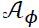 is trained on 100 media by minimizing the discrepancy between predicted and measured growth rates while enforcing flux non-negativity and stoichiometric constraints (see loss terms). Here, *S* is the stoichiometric matrix of the genome-scale metabolic model iML1515 (E. coli K-12 MG1655), 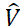 is the vector of predicted reaction fluxes of iML1515, and μ (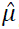) is the measured (predicted) growth rate. The AMN training is performed prior to solving ML tasks; consequently, the same optimal parameters ϕ^∗^and thus the same frozen operator *g*(.) are used across all ML problems in this study.

Our features-to-medium mapper is an MLP that selects one medium from a library of 280 experimentally prepared media, avoiding the need to design and prepare new media for each new ML task. This library was constructed as a diverse yet finite set of experimentally realizable growth conditions: all media share a minimal medium M9 base and vary by defined combinations of 3 carbon sources and 25 selected amino acids/nucleobases (Materials and Methods **M6.1**), yielding 280 distinct media that span a broad range of growth phenotypes. The library size presents a practical compromise between phenotypic coverage and experimental throughput: large enough to cover diverse nutrient regimes, yet small enough to prepare once as a reusable fixed library for many ML tasks.

To guide selection, the features-to-medium mapper relies on an *in-silico* predictor, an Artificial Metabolic Network (AMN), which estimates the strain’s growth rate for each candidate medium (**Fig. 3b**). During mapper training, the AMN provides a differentiable *proxy* signal (the predicted growth rate) to optimize medium selection. After training the mapper, *E. coli* (the reservoir) is grown on the selected medium, and the ML task prediction is produced from the measured reservoir response.

AMNs are neural–mechanistic hybrid models (Materials and Methods **M7**) that predict growth rate for a given medium and have recently been developed to improve the performance of flux balance analysis (FBA)^20^. Importantly, in the present study, the AMN is trained only once on 100 media and is not retrained for individual ML tasks; instead, it is frozen after training and reused as a common growth-rate predictor throughout the study (**Fig. 3c**). The 100-medium training set was drawn randomly from the 280-medium library. *E. coli* was grown on these media (Materials and Methods **M6.2**), and growth rates were measured (Materials and Methods **M6.3**). We observed that this 100-medium training set spans growth rates from 0.15 to 0.90 h^−1^ (cf. Supplementary Fig. **S6**), covering slow-growth to near-maximal growth regimes reported for *E. coli* K-12 MG1655 in glucose minimal media^21^.

Once a medium is selected within the 280-medium library, we grow the strain, and record the growth curve with a plate reader (Materials and Methods **M6.2**). We propose two different RC readouts to predict the ML problem output, one takes as input a time-series (the growth curve as in **Fig. 1.b**) and the other a scalar (the growth rate as in **Fig. 1.c**). With the growth curve, the readout is a linear classifier or regressor that takes as input 27 OD_600_ measurements over a time-series (from 0 to 16 hours) and produce the problem output. For the growth-rate readout, we normalize the measured growth rate and use it as an intrinsic scalar output of the reservoir; the task prediction is obtained directly from this value, without training a readout (additional information on the two readouts can be found in Materials and Methods **M8**).

### Solving regression and classification tasks with an *E. coli* reservoir

Throughout this section, we solve ML classification and regression tasks using the *E. coli* RC architectures shown in **Fig. 1b-c**, and we compare the results with standard classifiers and regressors using accuracies and regression coefficients computed by cross-validation, as detailed in Materials and Methods **M9**.

We first benchmarked our *E. coli* reservoirs on a set of linear and nonlinear classification tasks, similar to those studied by Baltussen *et al*.^11^. We considered eight two-dimensional classification problems, generating 500 randomly sampled input coordinates for each task (Materials and Methods **M10**). We used the features-to-medium mapper described above to select suitable media for these inputs. We then grew *E. coli* in the selected media, recorded growth curves, and computed growth rates; these measurements were used as reservoir states and fed to the readout (**Fig. 1b-c**).

We evaluated two readouts: one using the growth rate (scalar readout) and one using the full growth curve (time-series readout of 27 OD600 measurements). For the time-series readout, we used a linear classifier to predict the task label (*cf*. hyperparameters in Supplementary Table **S4**). We compared both readouts to standard classifiers: support vector machines (SVM), multilayer perceptrons (MLP), and XGBoost (XGB). As shown in **Fig. 4**, our *E. coli* RC consistently outperformed SVM and MLP and achieved performance close to XGB and the Formose chemical reservoir^11^.

**Fig. 4.**
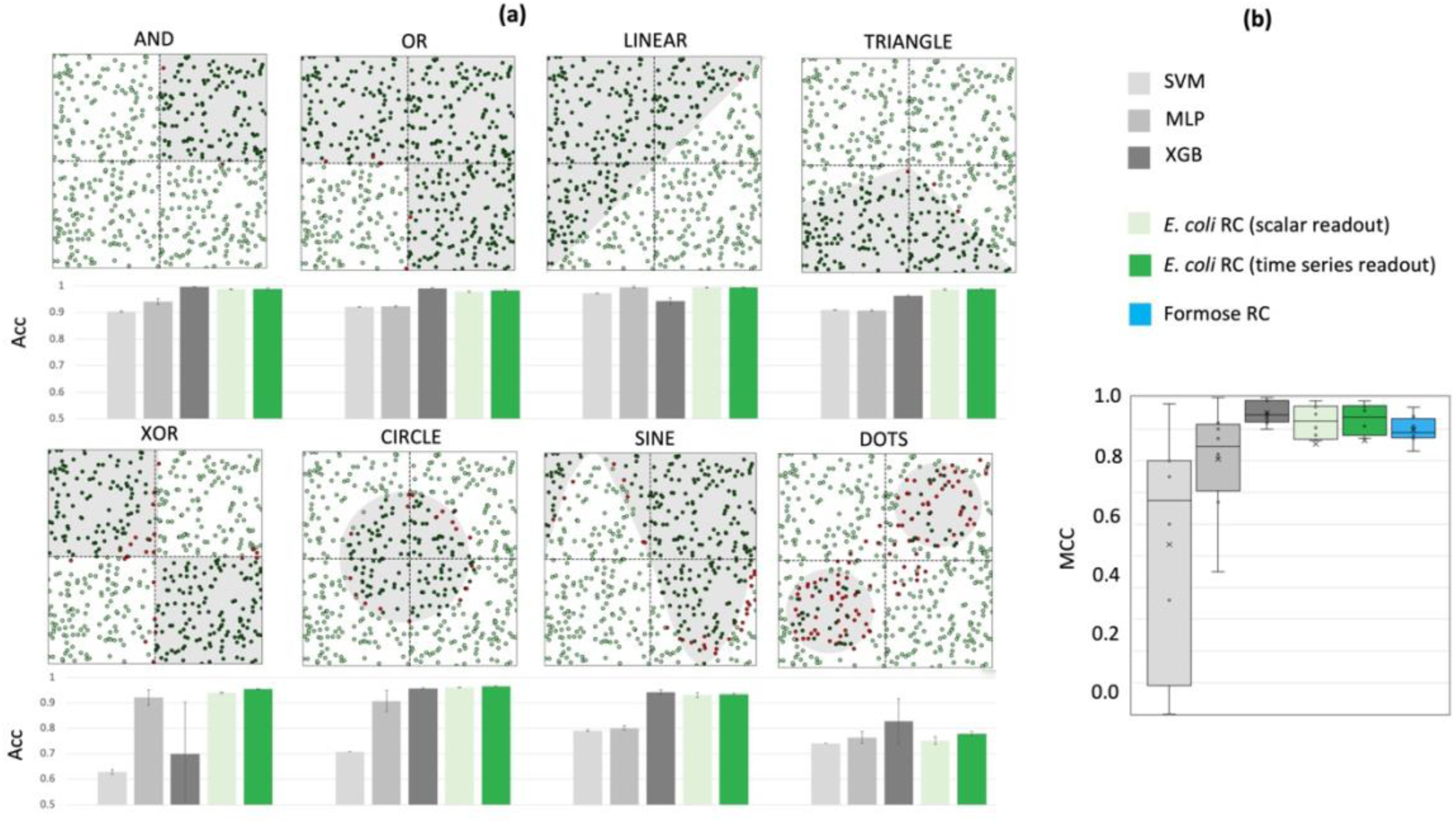
Classification problem performances with E. coli RC. **(a)** Performances for 4 linear (AND, OR, LINEAR, TRIANGLE) and 4 nonlinear (XOR, CIRCLE, SINE, DOTS) classification tasks with SVM (light gray), MLP (medium gray) and XGB (dark gray) classical classifiers along with E. coli RC systems based on growth rates (scalar readout, light green, architecture shown in Fig. 1c) and growth curves (time-series readout, dark green, architecture shown in Fig. 1b). Patterns are drawn following Materials and Methods **M10**. Red dots are false positives and false negatives. All accuracies are computed for 5-fold cross-validation (Materials and Methods **M9**). **(b)** Box plot for the 8 classification problems of panel a using Matthews Correlation Coefficient (MCCs) as a performance metric. As in Baltussen et al.^11^, MCC is calculated for 26-fold cross validation repeated 20 times. The data for the Formose Chemical Reservoir box-plot was extracted from Baltussen et al. other data are in the Supplementary file ‘Reservoir_Classification’.

We next applied our *E. coli* RC to 10 benchmark regression tasks from the OpenML platform (Materials and Methods **M11**). As for classification, the features-to-medium mapper transformed each input vector into a medium used to grow the *E. coli* strain. We again used two readouts: one using the growth rate (scalar readout) and one using the full growth curve (time-series readout of 27 OD600 measurements). For the time-series readout a linear regressor was used to predict the target value (*cf*. hyperparameters in Supplementary Table **S4**). The results in **Fig. 5** show that the scalar readout outperforms multi-linear regression (MLR) in most cases, while the time-series readout outperforms MLR and MLP and achieves performance comparable to XGB for mean and median values.

**Fig. 5.**
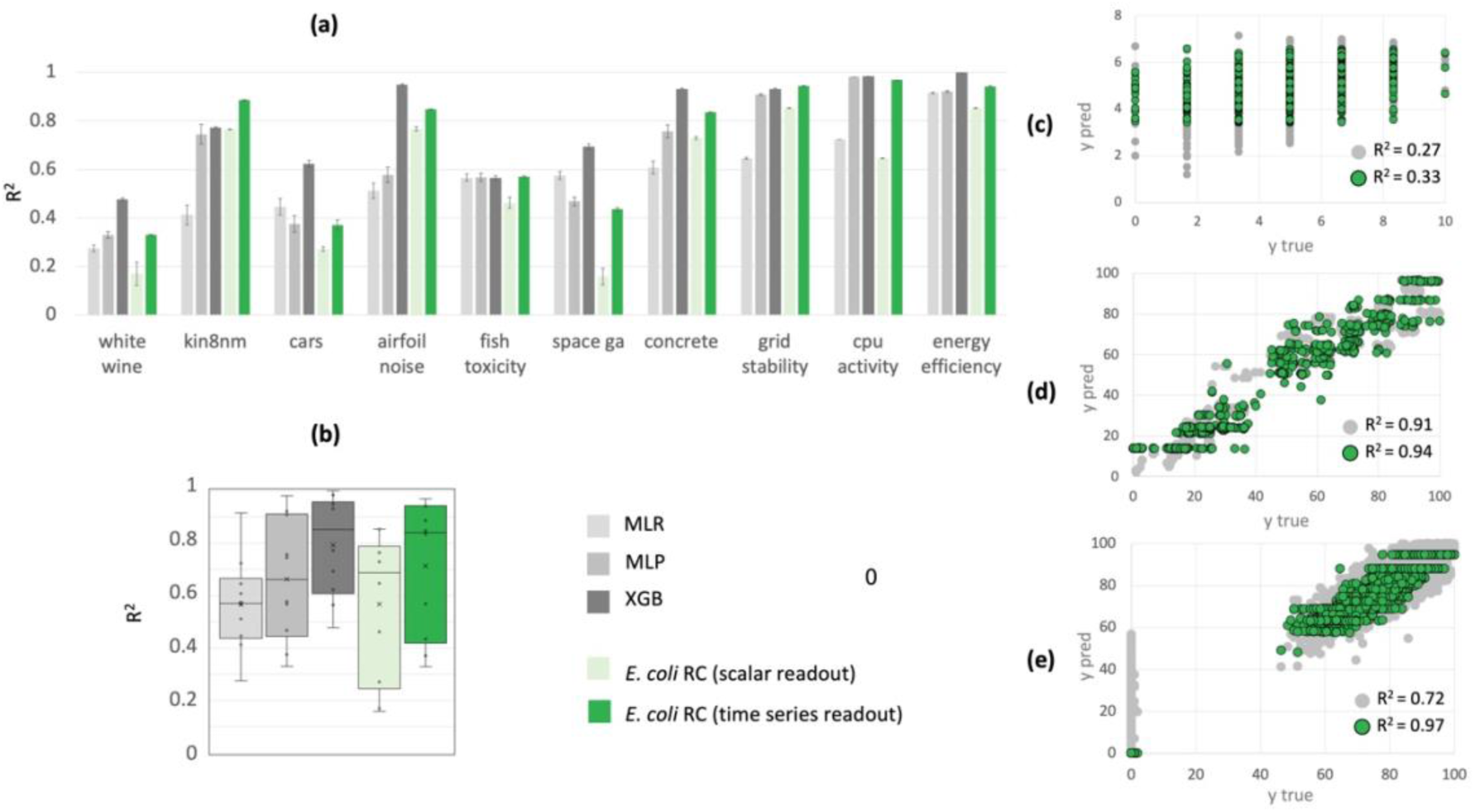
Regression problem performances with E. coli RC. **(a)** Performances for 10 regression tasks with MLR (light gray), MLP (medium gray) and XGB (dark gray) classical regressors along with E. coli RC systems based on growth rates (scalar readout, light green, architecture shown in Fig. 1c) and growth curves (time-series readout, dark green, architecture shown in Fig. 1b). The 10 regressions tasks were extracted from the OpenML platform (Method section **M11**). All R^2^ values are computed for 5-fold cross-validation (Materials and Methods **M9**). **(b)** Box plot for the same 10 regression tasks. **(c)** Results obtained for the ‘white wine’ regression problem with MLR (light gray) and E. coli RC based on growth curves (dark green). **(d)** MLR and E. coli RC results obtained for the ‘energy efficiency’ regression problem. **(e)** MLR and E. coli RC results obtained for the ‘cpu activity’ regression problem.

As already mentioned, a practical challenge in physical RC is how to present task inputs to the reservoir. In our framework, each input feature vector is encoded into an experimental condition, namely a growth medium selected from a fixed library. This encoding is performed by a trainable features-to-medium mapper (**Fig. 3a**), which select one medium per input among 280 experimentally prepared media.

Because the mapper is implemented as an MLP, this input-encoding step is non-linear. However, its role is limited to selecting an experimental medium, while the reservoir states arise solely from the strain’s growth response. Consistent with this, the time-series *E. coli* RC pipeline (inputs → mapper → medium → *E. coli* growth curve → linear readout) achieves higher performance than an MLP trained directly on the raw inputs: median MCC increases from ∼0.84 (MLP on inputs) to ∼0.93 (*E. coli* RC time-series), and mean R^2^ increases from ∼0.66 to ∼0.84 (**Fig. 4b and Fig. 5b**). These results indicate that coupling the same inputs to a living bacterial reservoir yields representations that are more informative for downstream prediction than a standard MLP baseline.

### Benchmarking GEM-based reservoirs

As we found that *E. coli* can serve as a reservoir to solve classification and regression tasks with performances comparable to, or better than, standard classifiers and regressors, we asked whether other microbial species could achieve similar results.

To address this question, we built *in-silico* reservoirs for 10 microbial species using their genome-scale metabolic models (GEMs). For each GEM, following Materials and Methods **M12**, we generated 10,000 media and computed growth rates by flux balance analysis (FBA)^20^. As illustrated in **Fig. 6a**, we used the features-to-medium mapper to convert the inputs of the 10 regression problems of **Fig. 5** into growth media, selecting one medium per input from the 10,000-medium library. The selected medium was then used as input to FBA, which produced a growth rate that was fed to the readout. Because FBA does not produce growth curves, we did not use a time-series readout. Instead, the normalized growth rate served as the reservoir state, and was used directly to predict the problem output (Materials and Methods **M8**).

**Fig. 6.**
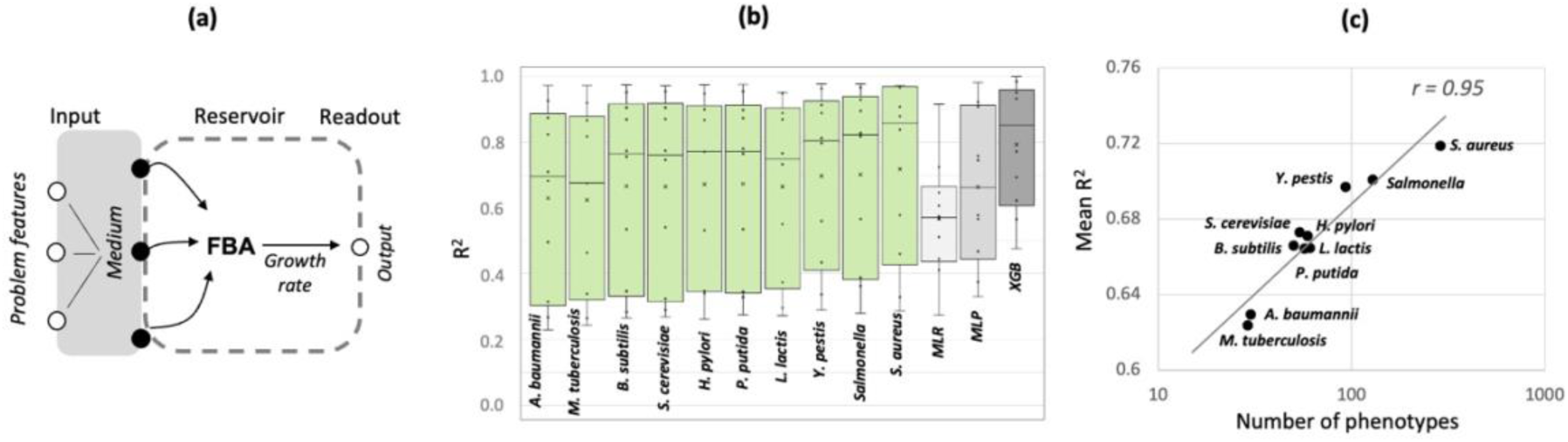
Regression problem performances with GEM RC. **(a)** Schematic of the RC system architecture. The architecture is similar to the one shown in Fig. 1c but the growth rate is calculated by FBA^20^ instead of being measured. **(b)** Box plots for regression coefficients obtained for 10 regression problems (Materials and Methods **M11**) and for 10 GEM RC systems (Materials and Methods **M12**). The GEM RC results are compared with those obtained using MLR, MLP, and XGB. **(c)** Mean regression coefficient (R^2^) vs. number of phenotypes obtained for the 10 GEMs of panel b. The Pearson correlation coefficient (r) between mean R^2^ values and the linear fit (y = 0.1 x + 0.5) is 0.95. All R^2^ values were acquired on 5-fold cross-validation sets for three repeats. Data are in the supplementary file ‘Reservoir_Regression’.

We compared the performance of these GEM-based reservoirs to MLR, MLP, and XGB. As shown in **Fig. 6b**, based on mean and median values, GEM-based reservoirs generally outperform MLR and MLP, and some GEM-based reservoirs achieve performance comparable to XGB. Performance also appears higher than in **Fig. 5b** with the scalar readout, most likely because the GEM analysis uses a much larger media library (10,000 media here versus 280 in **Fig. 5**). Most importantly, performance varies substantially across species.

To explore the performance differences between species, we quantified the number of distinct phenotypes each GEM can produce using the 10,000-medium library (Materials and Methods **M13**). Here, phenotypes were defined based on growth rate; accordingly, we computed the FBA calculated growth rate for each model and each 10,000 media. **Fig. 6c** shows a strong positive correlation between phenotypic diversity and regression performance. This suggests that strains producing only a few phenotypes (e.g., growth *vs*. no growth) are less suited to complex regression tasks. This may also explain why our *E. coli* physical reservoir performs better on classification tasks (**Fig. 4**) than on regression tasks (**Fig. 5**) when using the growth-rate readout, since binary classification requires only two separable responses.

## Discussion

This study demonstrates the potential of a physical *E. coli* reservoir, to address both regression and classification tasks.

The main key advantage of the *E. coli* reservoir compared to other published physical reservoirs ^11,12^ lies in its applicability to clinically relevant settings. Standard ML pipelines applied to untargeted metabolomics data on plasma samples yield accuracies around 0.9 for COVID-19 disease severity classification. Using growth curves, our *E. coli* reservoir distinguished between mild *vs*. severe COVID-19 cases with a cross-validated accuracy of 0.86 (AUC 0.90) with 20-fold cross-validation, and achieved even higher accuracy (0.96) and AUC (0.97) when classifying between COVID-19 positive and negative samples.

Our observed differences in OD profiles across sample categories are consistent with metabolite levels reported in the literature. Indeed, viral replication, organ dysfunction and immune cell expansion following a COVID-19 infection are responsible for wide metabolomic alterations in serum and plasma. These variations might explain the growth pattern differences observed in the different cases. Decreased levels of lipids and amino acids derivatives^13,23^ (mostly from the arginine metabolism) have been observed in COVID-19 patients compared to uninfected individuals^13^. Given that *E. coli* can catabolize some classes of lipids, these observations might explain the reduced OD values during the exponential and stationary phase when *E. coli* is cultured in plasma from COVID-19 patients. It should also be noted that increased levels of other carbon sources such as glucose, glucuronate or pyruvate have been reported in the plasma of COVID-19 patients^13,24^. However, it appears that the increased levels of these carbon sources doesn’t compensate for the effects of lipid depletion.

Regarding the mild vs. severe classification task, the more subtle differences in growth pattern may reflect more diverse alterations in the plasma metabolome at early stages of the infection. Decreased levels of several lysophosphatidylcholine (LCP) lipids and increased levels of kynurenine have been identified as negative prognosis biomarkers across a majority of the studies^13^. Altered levels of amino acids derivatives, nucleobases derivatives and polyamines have also been used as prognosis markers, though with less consistency across studies^14,16,25^. In a broader context, Maeda *et al*.^26^ demonstrated that amino acid deamination is associated with poor prognosis. Therefore, increased plasmatic levels of amino acid catabolites are expected for patients who will develop a severe form of the disease. Taken together, these variations may represent a plausible mechanistic basis for the growth differences observed in the mild vs. severe classification task.

Aside from using omics analyses only few studies have been carried out to predict COVID-19 severity and these generally include human health records as features^27^. Compared to these methods, our *E. coli* reservoir approach offers several advantages. Omics-based pipelines are generally expensive, time-consuming, and require specialized infrastructure and expertise. Our reservoir system achieves competitive accuracy without including patient data, operates with minimal cost and instrumentation, and produces results within hours using only a plate reader and bacterial cultures. These properties make it a highly accessible, scalable, and infrastructure-independent alternative for severity prediction, particularly valuable in decentralized or low-resource healthcare settings.

When applied to benchmark machine-learning tasks, our *E. coli* RC system outperforms standard classification and regression methods, including support vector machines (SVM), multi-linear regression (MLR), and multilayer perceptrons (MLP), as reflected by the mean and median values in the box plots in **Figs. 4-5**. Although the features-to-medium mapper is implemented as an MLP and therefore performs a nonlinear input encoding, its role is limited to selecting an experimental medium from a fixed library. The reservoir itself remains untrained; importantly, the full physical RC pipeline with time-series readout outperforms an MLP trained directly on the raw inputs (*cf.* box plots in **Figs. 4-5**), indicating that bacterial growth dynamics provide additional task-relevant computation beyond the encoder alone.

The improved performance observed for classification compared to regression likely reflects differences in task complexity: binary classification requires only two separable outputs, while regression demands broader phenotypic diversity. Supporting this, **Fig. 6c** shows a strong positive correlation between the number of internal states (phenotypes) and regression performances. This trend aligns with findings in reservoir computing, where increased state diversity enhances computational capacity^28,29,30^.

In our framework, an AMN model based on the *E. coli* GEM iML1515 is used only as a frozen guide for medium selection; downstream prediction is performed by the measured bacterial reservoir dynamics and the readout. Although AMN growth-rate prediction is not perfect (cf. Supplementary Fig. **S6**), an enzyme-constrained GEM (ecGEM^31^) could potentially refine medium ranking. However, using an ecGEM would require an AMN formulation compatible with enzyme-constrained models, which is not currently available but could be explored in future work.

Compared to chemical reservoirs^11^ and cortical neurons reservoirs (mBNNs)^12^, the *E. coli* reservoir offers a practical balance of classification performance, simplicity, and cost-effectiveness. The setup requires only standard microbiology equipment: an incubator and a plate reader with minimal consumables. In contrast, the chemical reservoirs depend on mass spectrometry and flow control systems, while mBNNs require optogenetics, microfluidics, and live-cell imaging, all of which demand substantial infrastructure and technical expertise. Despite this simplicity, the *E. coli* reservoir delivers competitive accuracies across a range of regression and classification tasks, relying solely on growth curves or growth rates as inputs to the readout.

All together this work supports the idea that living systems possess intrinsic computational capacity, echoing the concept of cellular computing supremacy proposed by Goñi-Moreno^32^. Our findings position bacterial RC as a biomimetic and infrastructure-light alternative capable of performing meaningful and useful information processing without the need for complex genetic re-engineering.

## Materials and Methods

### M1. Abbreviations

**AMN**: Artificial Metabolic Network, ***E*. *coli***: *Escherichia coli*, **FBA**: Flux Balance Analysis, **GEM**: Genome-scale metabolic model, **MLP**: Multi-Layer Perceptron, **MLR**: Multi-Linear Regression, **MS**: mass spectrometry, **OD**: Optical Density, **RC**: Reservoir Computing, **R**^2^: Regression coefficient. **SVM**: Support Vector Machine. **XGB**: XGBoost (parallel tree boosting).

### M2. COVID-19 patient plasma samples

All samples were extracted from the BIOMARCOVID cohort, a single-center, retrospective cohort study of patients hospitalized at the Grenoble-Alpes University Hospital (CHUGA) for COVID-19. This non-interventional, monocentric, retrospective study involving data and samples from human participants has been carried out in CHUGA according to French current regulation. The primary aim for that cohort was to develop a regression model that integrated clinical and biochemical parameters related to hospitalization and disease severity. The secondary aim was to identify differential metabolites and metabolic pathways between mild and severe COVID-19 outcomes using metabolomics analysis. Inclusion criteria included hospitalization for suspected COVID-19 and a positive PCR test for SARS-CoV-2. Exclusion criteria included age under 18, transfer from another hospital, opposition to research, ICU admission on the first day, and blood sample collection more than 48 hours after hospitalization at CHUGA. Clinical evaluation recorded demographic data (age, sex, BMI), disease progression (ICU, length of stay), and outcome status (class: mild or severe). Specialized teams assisted in acquiring clinical data. Patient outcomes were classified retrospectively using the WHO ten-category ordinal scale into six groups. Category 3: Patients hospitalized with No Oxygen supplementation; Cat 4: O2<=2L/min; Cat 5: O2>2L/min; Cat 6: admitted to ICU. The categories 3 and 4 were qualified as Mild group and 5 and 6 as severe group.

Anonymized patient data are is in the Supplementary files ‘Data_Covid_MS_ML.xls’ and ‘Reservoir_Covid.xls’. Subjects were all informed and did not oppose, written consent for participation was not required for this study in accordance with the national legislation and the institutional requirements. The study is registered on the Health Data Hub website under the number F20210218154851. A commitment to comply with Reference Methodology n°004 issued by French Authorities (CNIL) has been signed by the investigator (Prof. O. Epaulard co-author of the current study). Additional raw data regarding the cohort can be made available by the authors of the current study within respect of General Data Protection Regulation, without undue reservation.

### M3. Growth experiments on patients’ plasma samples

Nutritionally speaking, human plasma fulfills all growth requirements of *E. coli* and belongs to the rich media category. It contains magnesium ions, trace elements, several nitrogen, sulfur and phosphate sources^33^. It also contains all amino acids and some derivatives (citrulline and ornithine)^34^, several nucleobases and derivatives as well as vitamins, sugars, organic acids and free fatty acids^33^. Besides, plasmatic pH is around 7.4 and plasmatic osmolarity oscillates between 285 and 307 mOsm/kg in physiogical states^35^. Both parameters have optimal values for *E. coli* ^25,36^.

However, human plasma is not a suitable bacterial culture medium. The complement system prevents bacterial growth by lysing bacterial cells^37^. Complement inactivation strategies include heat inactivation and plasma deproteinization. Here, plasma deproteinization was preferred to protect heat-sensitive plasma components and because removing all plasmatic proteins will improve the accuracy of optical density measurements at 600nm.

Classical plasma deproteinization techniques used in analytical chemistry include usage of organic solvants ^38^ toxic for *E. coli* and were therefore dismissed in favor of filtration with a cutoff of 3kDa.

#### M3.1. Plasma deproteinization

The deproteinization process enables to inactivate the supplement system by removing its proteins from the plasma samples. Samples were centrifuged (1h, 4000g) in Amicon Ultra-4 Centrifugal Filters (3 kDa MWCO, 4 mL of sample volume), resuspended and re-centrifuged (1h, 4000g) in the same centrifugal filters. Filtrates were transferred and aliquoted into 1mL sterile tubes (100µL of deproteinized plasma per tube). The plasma selected for this series of experiments came from patients that were following antibiotic treatment before the blood sampling. The absence of antibiotics in the plasma samples was verified during the growth monitoring of the physical E. coli reservoir on plasma samples (presence of antibiotics should result in the absence of bacterial growth or an elongated lag phase). This procedure confirmed that no patients’ plasma used in this study contained antibiotics.

#### M3.2. Supplementation of magnesium ions

Another limitation arises from the blood collection technique used in this study. Indeed, blood samples were collected in tubes pre-filled with ethylenediaminetetraacetic acid (EDTA), a commonly used anticoagulant. EDTA chelates divalent cations such as magnesium ions^39^ which are critical for *E. coli*’s growth. *E. coli* was grown in deproteinized plasma samples from COVID-19 negative patients supplemented with a range of concentrations of magnesium and other (zinc, manganese, iron and copper) ions in order to assess the quantity to add to counterbalance the EDTA effects and ensure optimal growth. Optimal growth was defined using 2 parameters: (i) an increased growth rate and (ii) an increased maximal biomass (natural logarithmic transformation of the OD600nm measured at the beginning of the stationary phase). Supplementary Figure **S5** demonstrates that both criteria are respected when at least 0.82 µmol of magnesium sulfate is supplemented. The effect of other divalent cations is negligible in this case. It is worth noting that if less than 0.61 µmol of magnesium sulfate is supplemented, supplementation of other trace elements seems beneficial for *E. coli*’s growth. It might be explained by dilution effect: as the concentration of divalent cations increases while the amount of EDTA remains constant, a greater proportion of magnesium ions becomes unchelated. This effect may be accentuated by the fact that Zn^2+^, Cu^2+^, Fe^2+^, Ca^2+^ (naturally present in plasma) and Mn^2+^ have a higher affinity for EDTA than Mg^2+ 39^.Plasma samples were therefore supplemented with 1.02 µmol of MgSO_4_ throughout this study.

#### M3.3. Supplementation of a phosphate buffer

In physiological states, blood pH is buffered around 7.4 by the bicarbonate buffer system (HCO_3_^−^/H_2_CO_3_). This system is maintained by two organs: (i) the lungs that excrete CO_2_ through respiration and (ii) the kidneys which regulate the concentration of bicarbonate ions ^40^. Therefore, this system is ineffective in plasma samples were the concentrations of both CO_2_ and bicarbonate ions are not regulated anymore. Plasmatic proteins also have buffering capacity ^41^ but have been removed from the samples to inactivate the complement system. In order to buffer the pH during bacterial growth, a phosphate buffer (HPO_4_^−^ /H_2_PO_4_^2−^) is supplemented to the plasma samples.

#### M3.4. Growth monitoring of the E. coli reservoir on plasma samples

Growth curves are sensitive to experimental conditions (e.g., medium composition, temperature, shaking, and inoculum), particularly in lag time and overall trajectory. As shown next, we minimized variability by using a standardized protocol and processing samples/replicates in the same batch, resulting in highly reproducible growth curves across biological duplicates (average standard deviation of 0.4% across OD measurements).

*E. coli* K-12 MG1655 was cultured in all deproteinated patients’ plasma samples (37 mild, 36 severe and 27 negative) in order to assess its ability to classify: (i) plasma samples from early symptomatic mild vs. severe patients and (ii) COVID-19 infected patients vs. negative blood donors.

A glycerol stock of *E. coli* was streaked onto a M9 agar plate supplemented with 3.89 mM (0.07%) D-(+)-glucose (to match a physiological glycemia) and 0.5% yeast extract which was then incubated overnight at 37°C. A colony was used to inoculate a preculture of 3 mL of M9 liquid medium supplemented with 3.89 mM (0.07%) D-(+)-glucose and 0.5% yeast extract in a 14 mL preculture tube. The tube was incubated at 37°C under agitation at 180 rpm (Multitron, Infors HT) until exponential phase with an optical density at 600 nm measured between 0.5 and 1 (UVisco V-1100D spectrophotometer, LC-Instrument).

The plasma samples used in this study were collected in tubes containing ethylenediamine-tetra-acetic acid (EDTA), a commonly used anticoagulant to prevent blood clotting. EDTA chelates divalent cations such as MgSO_4_ which is required for *E. coli*’s growth. Moreover, the bicarbonate buffer system naturally present in the plasma is ineffective here. In light of these constraints, optimal quantities of MgSO₄ and a phosphate buffer were added to the samples before inoculation in order to ensure optimal *E. coli* growth.

In each well of a clear flat-bottom 96-well plate (except the edge wells that were filled with sterile milliQ water to avoid the edge effect), 1.02 µmol of MgSO_4_, 4.78 µmol of Na_2_HPO_4_ and 2.2 µmol of KHPO_4_ were added to 50 µL of deproteinized plasma. Sterile milliQ water was added to reach 100µL. Each well was inoculated with the *E. coli* preculture so that the starting optical density at 600 nm was estimated at 0.005. The plate was then incubated in a plate reader (BioTek HTX Synergy, Agilent Technologies) at 37°C with continuous orbital shaking at 807cpm (1mm). Bacterial growth was assessed by monitoring the optical density at 600 nm every 10 minutes for a period ranging from 8 to 24 hours. A technical duplicate was conducted for each sample by repeating the full procedure.

### M4. Statistical Modeling of *E. coli* growth curves on patients’ plasma samples

We investigated whether there were significant differences in *E. coli* growth dynamics between two independent group comparisons: negative *vs* positive, and mild *vs* severe. For each plasma sample, optical density (OD) measurements were recorded over time, with two technical replicates per time point (*cf.* Materials and Methods **M3**). To reduce measurement noise, and because replicate-specific effects were not of interest, we used the average OD across replicates at each time point for further analysis.

To evaluate group-specific differences in growth patterns, we fitted Generalized Additive Mixed Models (GAMMs)^17^ with the following structure:

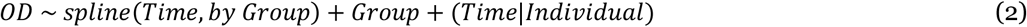

in this model:

– *spline* (*Time, by Group*) specifies group-specific smooth functions of time, allowing for different temporal trajectories between groups.
– *Group* is included as a fixed effect to test for differences in average OD levels between groups.
– (*Time|Individual*) represents a random effect accounting for between-individual variability over time.

A significant *p*-value (*p* < 0.05) for either the fixed group term or the group-specific smooth term indicates a meaningful group effect—either in the overall OD level or in the shape of its progression over time.

To further improve model fit and account for serial correlation in the time-series data, an autocorrelation correction was applied, which significantly reduced residual variation (*p* = 0.01). The first two hours of measurements were excluded from the analysis, as OD values during this period were too low to hold biological relevance. All analyses were conducted using R (version 4.4.3).

Results are presented in Supplementary Table **S3**. For negative and positive cases, GAMMs found statistically significant differences between groups in both the average OD across time points (*p*-value = 5.37 × 10⁻⁵) as well as in the temporal trajectories of OD over time (*p*-value < 2 × 10⁻¹⁶). For mild vs. severe cases, the overall mean OD did not differ significantly between the groups (*p*-value = 0.654). However, GAMMs revealed the presence of significant group-specific smooth terms (p-value < 2 × 10⁻¹⁶) indicating that the temporal evolution of OD differed substantially between mild and severe cases. These findings imply that the dynamics of how OD changes over time are distinct between the mild and severe groups, as in severe case, growth is delayed compare to mild cases: the mild mean OD was higher than severe during the first 4 hours and lower than severe after 4 hours.

### M5. Features-to-medium mapper to solve ML problems

The features-to-medium mapper is essentially an MLP with an input layer size equal to the number of features in the machine-learning problem to be solved and an output layer size equal to the number of media library (280). The MLP consists of three hidden layers, each of size 280 (*cf.* hyperparameters in Supplementary Table **S4**). To select one medium among the 280, we could apply a softmax activation function, which yields the probability of selecting each medium. However, we need to transform this probability vector into a one-hot vector, where the highest probability is set to 1 and all others to 0. To achieve this one-hot encoding, we use Gumbel–Softmax^22^, a differentiable approximation to argmax. Gumbel–Softmax uses a temperature parameter; during training, we start with a high temperature and gradually reduce it at each epoch. We found that using a logarithmic temperature decay provides better results than a linear decay.

As shown in **Fig. 3**, the loss function of the features-to-medium mapper relies on the predicted growth rate calculated for the selected medium. This loss function 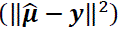 enforces the normalized predicted growth rate 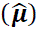 to match the normalized problem output (***y***). The predicted growth rate is obtained from the operator 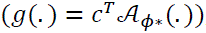, where *c*^*T*^extracts the growth rate (biomass flux) from the predicted flux vector (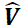). The flux vector 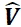 is computed by a trained AMN 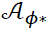 with optimal parameters ϕ^∗^. The AMN training procedure is detailed in section **M7**.

When training the features-to-medium mapper, the number of epochs was set to 1000 and an adaptive learning rate starting at 10^−3^ was used with the Adam optimizer^42^. Cross-validation (*cf.* section **M9**) was systematically used for all runs reported in **Figs. 4–6**. Batch sizes were set to 16 for training set comprising fewer than 2000 elements, and to 128 otherwise. All runs were carried out 10 times to produce different media fed to the *E. coli* reservoir (*cf*. Supplementary Table **S4** for additional hyperparameters).

### M6. *E. coli* physical reservoir to solve ML problems

We provide in this section the methods used to prepare culture media, to monitor growth of *E. coli* K-12 MG1655 (accession number: NC_000913.3), and to compute growth rates. The steps below were used to generate the 100-medium training set to train the AMN used in the features-to-medium mapper, and to prepare the 280-medium library used to produce growth curves

#### M6.1. Culture media preparation

All the media were based on a M9 minimal medium composed of (i) M9 salts at pH = 7.2: 47.76 mM Na_2_HPO_4_ (Merck/Sigma-Aldrich, CAS number: 7558-79-4), 22.04 mM KH_2_PO_4_ (Merck/Sigma-Aldrich, CAS number: 7778-77-0), 8.56 mM NaCl (Merck/Sigma-Aldrich, CAS number: 7647-14-5), 18.7 mM NH_4_Cl (Merck/Sigma-Aldrich, CAS number: 12125-02-09), (ii) magnesium sulfate: 4.08 mM MgSO_4_ (Merck/Sigma-Aldrich, CAS number: 7488-88-9), (iii) calcium chloride dihydrate: 0.1 mM CaCl_2_·2H_2_0 (Merck/Sigma-Aldrich, CAS number: 10035-04-08), and (iv) trace elements: 0.4 mM Na_2_EDTA·2H_2_O (Merck/Sigma-Aldrich, CAS number: 6381-92-6), 0.03 mM FeCl_3_·6H_2_O (Merck/Sigma-Aldrich, CAS number: 7705-08-0), 6.16 µM ZnCl_2_ (Merck/Sigma-Aldrich, CAS number: 7646-85-7), 0.76 µM CuCl_2_·2H_2_O (Merck/Sigma-Aldrich, CAS number: 7447-39-4), 0.42 µM CoCl_2_·6H_2_O (Merck/Sigma-Aldrich, CAS number: 7791-13-1), 1.62 µM H_3_BO_3_ (Merck/Sigma-Aldrich, CAS number: 10043-35-3), 0.08 µM MnCl_2_·4H_2_O (Merck/Sigma-Aldrich, CAS number: 13446-34-9), 1.65 µM Na_2_MoO_4_·2H_2_O (Merck/Sigma-Aldrich, CAS number: 7631-95-0).

This minimal medium was supplemented with different combinations of nutrients selected from 3 sugars or acids, 20 proteinogenic amino acids and 5 nucleobases. The 3 sugars or acids used as carbon source and their final concentrations are: 44.4 mM D-(+)-glucose, 53.29 mM D-(+)-xylose (Merck/Sigma-Aldrich, CAS number: 58-86-6), 49.37 mM sodium succinate dibasic (Merck/Sigma-Aldrich, CAS number: 150-90-3). Final amino acids and nucleic bases concentrations used are based on the EZ rich defined medium composition^43^: 0.8 mM L-alanine (Merck/Sigma-Aldrich, CAS number: 56-41-7), 5.2 mM L-arginine (Merck/Sigma-Aldrich, CAS number: 74-79-3), 0.4 mM L-asparagine (Merck/Sigma-Aldrich, CAS number: 70-47-3), 0.4 mM L-aspartic acid (Merck/Sigma-Aldrich, CAS number: 56-84-8), 0.1 mM L-cysteine (Merck/Sigma-Aldrich, CAS number: 52-90-4), 0.55 mM L-glutamic acid (Merck/Sigma-Aldrich, CAS number: 56-86-0), 0.615 mM L-glutamine (Merck/Sigma-Aldrich, CAS number: 56-85-9), 0.8 mM glycine (Merck/Sigma-Aldrich, CAS number: 56-40-6), 0.2 mM L-histidine (Merck/Sigma-Aldrich, CAS number: 71-00-1), 0.4 mM L-isoleucine (Merck/Sigma-Aldrich, CAS number: 73-32-5), 0.8 mM L-leucine (Merck/Sigma-Aldrich, CAS number: 61-90-5), 0.5 mM L-lysine (Merck/Sigma-Aldrich, CAS number: 56-87-1), 0.2 mM L-methionine (Merck/Sigma-Aldrich, CAS number: 63-68-3), 0.4 mM L-phenylalanine (Merck/Sigma-Aldrich, CAS number: 63-91-2),0.4 mM L-proline (Merck/Sigma-Aldrich, CAS number: 147-85-3),10 mM L-serine (Merck/Millipore, CAS number: 56-45-1), 0.4 mM L-threonine (Merck/Sigma-Aldrich, CAS number: 72-19-5), 0.1 mM L-tryptophan (Merck/Sigma-Aldrich, CAS number: 73-22-3), 0.2 mM L-tyrosine (Merck/Sigma-Aldrich, CAS number: 60-18-4), 0.6 mM L-valine (Merck/Sigma-Aldrich, CAS number: 72-18-4), 0.2 mM adenine (Merck/Sigma-Aldrich, CAS number: 73-24-5), 0.2 mM cytosine (Merck/Sigma-Aldrich, CAS number: 71-30-7), 0.2 mM uracil (Merck/Sigma-Aldrich, CAS number: 66-22-8), 0.2 mM guanine (Merck/Sigma-Aldrich, CAS number: 73-40-5), 0.2 mM thymidine (Merck/Sigma-Aldrich, CAS number: 50-89-5).

The objective was to develop a set of media capable of supporting a broad spectrum of maximal growth rates for *E. coli* K-12 MG1655. To this effect, different combinations of amino acids, nucleobases, and other carbon sources were designed to supplement a M9 minimal medium. Amino acids involved in the same biosynthesis pathway were grouped together as well as purines and pyrimidines (regarding nucleobases). 9 groups of amino acids were formed: Gly/Ser/Thr, Asn/Asp/Ala, Glu/Gln, Arg/Pro, Val/Leu/Ile, Lys, Cys/Met, His, Phe/Tyr/Trp (His and Lys were often associated together for the sake of simplicity, Glu and Gln were separated from Asn/Asp/Ala to ensure balanced groups). Each group was ranked according to their metabolic cost for *E. coli*, as defined in Zampieri *et al.* ^44^. It has been demonstrated that amino acids with a low metabolic cost are consumed at a higher rate than the other ones ^44^. The same study suggested that these low-cost amino acids could be used as carbon and energy sources ^44^. Therefore, the hypothesis underlying the ranking was that amino acids with lower metabolic costs would have a greater contribution to the growth rate than the other ones. By varying the number of groups of amino acids supplemented in the medium as well as their rank, it was possible to generate a first set of media capable of supporting a wide range of maximal growth rates for *E. coli*. This range was further widened by designing new media supplemented with amino acids but also nucleobases and different energy-rich or energy-poor carbon sources (D-(+)-glucose, D-(+)-xylose, succinate).

#### M6.2. Bacterial growth monitoring

An *E. coli* K-12 MG1655 glycerol stock was streaked onto a M9 agar plate supplemented with 44.4 mM (0.8%) D-(+)-glucose (the composition is described in « Bacterial strain and culture media ») which was then incubated overnight at 37°C. A colony was used to inoculate a preculture of 10 mL of M9 liquid medium supplemented with 44.4 mM D-(+)-glucose in a 100 mL Erlenmeyer flask. The flask was incubated at 37°C under agitation at 180 rpm (Orbital floor shaker ZWY-B3222, Labwit) until the optical density at 600 nm of the culture reached 0.5 (measured on a UVisco V-1100D spectrophotometer, LC-Instru). 198 µL of each medium to test was prepared in each well of a clear flat-bottom 96-well plate (except the edge wells that were filled with sterile milliQ water to avoid the edge effect) and inoculated with the *E. coli* preculture so that the starting optical density at 600nm was estimated at 0.005. Each medium was tested in triplicate. The plate was then incubated in a plate reader (BioTek HTX Synergy, Agilent Technologies) at 37°C with continuous orbital shaking at 807cpm (1mm). Bacterial growth was assessed by monitoring the optical density at 600nm every 20 minutes for a period ranging from 16 to 24 hours.

#### M6.3. Growth rate determination

A natural logarithm function was applied to the optical density measurements at 600 nm. First, the three timepoints between which the growth rate is maximal were determined using linear regression. The growth rate determination window was then extended in order to cover all timepoints of the exponential phase. To do so, a fitting score was calculated between the linear model and the experimental data. The growth rate determination window was expanded in both directions (before and after the two initial timepoints) as long as the fitting score remained below a chosen threshold. The maximal growth rate was finally calculated using the timepoints within the final window. Each medium was tested in triplicate. The mean and standard deviation of each triplicate was computed. The training set file contained the media (in a binary form capturing the presence or absence of a specific medium metabolite) the average growth rate and the corresponding standard deviation. In addition to growth-rate we also computed lag time and maximal density (OD_MAX_). We define the lag time as the last time point for which the slope did not exceed 25% of the growth rate, and OD_MAX_ as the average of the top 5 highest OD values.

### M7. AMN hybrid models

AMNs^45^ are hybrid neural-mechanistic models which purpose is to surrogate Flux Balance Analysis (FBA)^20^ to better fit experimental measurements. FBA is a well-established technique in systems biology and bioengineering used to model the metabolic behavior of genome-scale metabolic models (GEMs)^46^. Among the various AMN architectures propose in Faure *et al*.^45^, we chose AMN-QP. The architecture of AMN-QP, shown in **Fig. 3c**, is inspired by Physics Informed Neural Networks (PINNs). As in PINNs, the loss function comprises three terms. The first 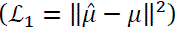 enforces growth rate predictions (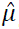) to match the measurements (μ). The second 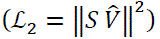 is the classical FBA problem constraint, *S V* = 0, where *S* is the stoichiometric matrix of the model (iML1515) corresponding to the strain (MG1655). The third loss 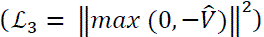 enforces the boundary condition where predicted fluxes (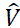) must remain positives. During training, all the terms are collected into one loss 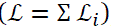, scaling is performed using the prior loss methods where 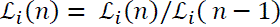 with *n* being the epoch number.

AMN was trained on both experimentally acquired growth rates and FBA calculated growth rates. As further detail in Supplementary Table **S4,** the AMN architecture was composed of an MLP with one hidden layer, with an input size of 28, a hidden layer of size 500 and an output vector (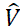) of a size equal to number of mono-directional reactions (cf. Supplementary Table **S5**). The model was trained for 1000 epochs, the train rate was set to 10^−3^ with Adam as the optimizer^42^.

The experimental training set used to train the AMN (within the features-to-medium mapper) for the *E. coli* reservoir experiments reported in **Figs. 4-5** consists of 100 media with measured growth rates (see section **M6** and Supplementary file “Data_Ecoli.xls”). Results for cross-validation with this AMN training set are provided in Supplementary Figure **S6**.

The AMN training sets used in **Fig. 6** were generated for 10 different GEMs. For each GEM, 10 000 media were created as detailed in section **M12** and FBA was run to compute the growth rate for each medium and GEM. In cross-validation, all AMN models achieved R² > 0.99 between predicted and FBA-computed growth rates, largely due to the large size of the training sets.

### M8. *E. coli* reservoir readouts to solve ML problems

We propose two different readouts. The first is using *E. coli* reservoir growth curves (as in **Fig. 1b**). Precisely for each ML problem instance and associated growth medium, all OD_600_ values taken along 47 times points of the growth curve are fed to a standard linear classifier or linear regressor depending on the ML problem to be solved. Both the classifier and the regressor are taken from SciKitLearn ^47^ and are making use of Ridge regularization (*cf.* hyperparameters in Supplementary Table **S4**). The readout results labeled ‘time-series ‘ shown in **Fig. 3** and **Fig. 4** were obtained for 5 –fold cross validation and three repeats and we report the top three accuracies (for classification) or regression coefficients (for regression) across different media selected by the features-to-medium mapper.

The second readout is making use of the growth rate calculated from growth curve for each ML problem instance and associated growth medium (as in **Fig. 1c**). Here we simply normalize both the measured growth rate and the problem readout and the problem readout can directly be predicted from the growth rate. The readout results labeled ‘scalar’ shown in in **Fig. 3** and **Fig. 4** were obtained for 5 –fold cross validation and three repeats and we report the top three accuracies (for classification) or regression coefficients (for regression) across different media selected by the features-to-medium mapper.

### M9. Cross-validation procedure

We used the same cross validation method for all regressors and classifiers on all training sets. The training set is randomly divided into *k* folds. One-fold (validation set) is selected to make prediction while training is performed on the remaining *k*-1 folds. The process is repeated selecting each fold as a validation set one after another. All regressions coefficients and accuracies reported in the manuscript and Supplementary Figures and Tables were computed using predicted values from the *k* aggregated validation sets. One notes that when the validation set contains only one element (*k* equals the size of the training set) our cross-validation method is Leave One Out (LOO).

To obtain statistics for accuracies and regression coefficients, unless specified otherwise, we repeated all cross-validation processes three times.

### M10. Classification problems

To generate data for the classification tasks shown in **Fig. 3** we draw pairs from a uniform [0,1] distribution to get 500 samples with independence x_1_ and x_2_ features. Similar to Baltussen *et al.*^11^, 8 different conditions (AND, OR, LINEAR, TRIANGLE, XOR, CIRCLE, SINE, DOTS) were used to calculate classes labeled 0 or 1. To produce **Fig. 3**, we used the SVM, MLP and XGB classifiers from SciKitLearn ^47^ (*cf.* Baseline comparator in Supplementary Table **S4**) and acquired accuracies on cross-validation sets (cf. section **M9**). We iterated three times to compute standard deviation

### M11. Regression problems

The 10 regression problems downloaded from OpenML are listed in Supplementary Table **S6.** To compare conventional and physical reservoirs regression results in **Fig. 4**, we used the MLR, MLP and XGB regressors from SciKitLearn^47^ (*cf.* Baseline comparator in Supplementary Table **S4**) and acquired regression coefficients (*R*^2^) on cross validation sets (cf. section **M9**). We iterated three times to compute standard deviation.

### M12. Genome Scale Metabolic Models (GEMs)

A set of 10 GEMs were downloaded from the BiGG database^46^, these are listed in Supplementary Table **S5**. Following a method outlined in Faure *et al*.^45^ metabolic reactions having no specific directions were duplicated one forward one backward. A set of 10,000 media were generated for each of the 10 GEM models. Each medium comprised two parts: a minimal medium and a variable medium. The minimal media were specific to each model, and defined as the set of metabolites necessary for the model to produce a growth. The variable media were generated at random among the same 28 metabolites listed in Materials and Methods **M6.1**. As with the experimental training set each variable medium was represented by a 28 bits vector. FBA was run using Cobrapy^48^ for each medium and each model, all reaction rates (including the growth rate) were collected and the growth rate was stored in the training set files.

### M13. Phenotype calculations

To compute the number of phenotypes in **Fig. 6**, for the 10 strains considered in this study we used the 10,000 instances training sets described in Materials and Methods **M12**. The number of different phenotypes is simply the number of different growth rates. Several media can lead to the same growth rate especially if one does not distinguish two rates when they are separated by a distance lower than a predefined threshold. We chose an arbitrary threshold of 0.01 mol h^−1^ cell^−1^ which is the limit of what can be measured experimentally using a spectrophotometer. We note that the other thresholds we tested (0.05, 0.005 and 0.001) produced trends similar to those in **Fig 6c**.

## Supporting information

Supplementary data files (zip)

## Acknowledgements

Authors would like to acknowledge funding provided by the ANR funding agency grant numbers ANR-22-PEBB-0008 (PEPR B-BEST France 2030 program) and ANR-21-CE45-0021-01 (AMN project) and the UE HORIZON BIOS program (grant number 101070281). J.L.F is getting support from all programs, A.L.G. is supported by the ANR-21-CE45-0021-01, the ANR-24-CE44-1190 project, the Foundation University Grenoble Alpes and Foundation Air Liquid (BIOMARCOVID Project), P.A. by the AMN project, A.H. by the BIOS project, P.M. by the PEPR B-BEST project. We thank Alexandra Martin (INRAE, University of Paris Saclay) for her help in acquiring *E. coli* growth curves for different media, and we thank Claire Roman (Campus Innovation Paris, Air Liquide & IRIMAS, University Haute Alsace) for her help and suggestions regarding the statistical analyses. We thank Laura Alonso and Xavier Hinaut (INRIA, U. Bordeaux) for reading our manuscript and providing helpful suggestions.

## Author Contributions

P.A. performed all microbiology work and generated all training sets used in the study growing *E. coli* K-12 MG1655 for different media and on COVID-19 samples. T.N.A.H. performed the statistical analysis. P.M. created and maintained the GitHub implementation and tested all codes. O.E. and A.L.G wrote the clinical trial. O.E. and S.B. collected the plasma samples. F.F., F.C., and A.L.G performed metabolomics experiments. J.L.F designed and supervised the study, acquired the fundings, and wrote the reservoir and machine learning codes. All authors contributed to writing manuscript sections, reading and editing the manuscript. All authors read and approved the manuscript.

## Data Availability

Aside from the Supplementary files linked to this manuscript three folders are used to run the code: a folder with data used as input to the code, a folder with saved trained models, and a folder with results. These are available at: https://doi.org/10.5281/zenodo.14961167

## Code Availability

The source code associated to the paper and the notebooks used to produce the results are available at: https://doi.org/10.5281/zenodo.15280670 as well at the following GitHub repository: https://github.com/brsynth/bacterial_rc

## Supplementary Information

Reservoir_Supplementary.pdf, for Supplementary Notes, Tables and Figures. Covid_Data_MS_ML.xls, Reservoir_Covid.xls, Ecoli_Data.xls, Reservoir_Regression.xls, Reservoir_Classification.xls, for the data used to run the code and to generate all Figures and Tables.

