## Supplementary Information for "Living Bacterial Reservoir Computers for Information Processing and Sensing"

**This PDF file includes:**

Methods MS1 to MS3
Tables S1 to S6
Figures S1 to S8
SI References

**Other supporting materials for this manuscript include the following:**

Datasets S1 to S5

### Supplementary Methods

#### MS1. Metabolomics analyses on plasma samples

Metabolomics analysis was carried out on 81 patient samples (50 mild, 31 severe) from the BIOMARCOVID cohort. Mass spectrometry (MS) data were acquired using two liquid chromatography (LC) columns C18 and HILIC as described below along with the methods we used for annotation. Metabolomics data is given in the Supplementary file 'Covid_Data_MS_ML'.

***LC-MS analysis with C18***. We used a UHPLC Vanquish Flex Binary system coupled with an Orbitrap Q-Exactive Plus mass spectrometer (Thermo Fisher Scientific). Chromatographic separation employed an Accucore RP-MS column with a gradient elution: phase A (water with 0.1% formic acid) and phase B (acetonitrile with 0.1% formic acid). Flow rate was set at 0.4 mL/min, injection volume at 5 µL, and column temperature at 30 °C. Mass spectrometry was conducted in positive ion mode (m/z 100–1500). Full-scan resolution was 70,000, with MS/MS at 17,500 and collision energies of 10, 35, and 55 eV. An exclusion list was applied based on blank sample analysis. Data (.raw) were converted to .mzXML using ProteoWizard and processed with XCMS (Benton *et al*, 2008) in R (https://github.com/sneumann/xcms).

Preprocessing of the .mzXML raw files to obtain the feature Table was carried out with R version 4.1.2, in Rstudio environment (version 1.4.1717), using XCMS package (version 3.16.1) as follows: Chromatographic peak detection was performed with centWave algorithm (peakwith = 10-30s, ppm = 10, snthresh = 6, mzdiff = 0.01); PeakGroupsParam class (minFraction =0.7,span=0.2) was used to perform the alignement step; chromatographic peaks correspondence from different samples was achieved by using the groupChromPeaks function (minFraction = 0.7, bw = 5); filling missing peaks was done using the ChromPeakAreaParam method with standard parameters. Features annotation from XCMS (isotopic features and adduct formations, dimers, multimers, neutral losses) was carried out using CAMERA package (Kuhl *et al*, 2012) (version 1.48.0) with parameters perfwhm = 0.6, mzabs = 0.01, cor_eic_th = 0.75.

Quality control on the processed data was assessed by plotting a PCA score plot and a run plot of intensities sum vs injection order. Then, filtering of processed data was performed by removing features with a coefficient of variation (CV) above 30%, calculated across QC samples. Missing values imputation was carried out by replacing missing values with 1/5 of the minimum positive value of each variable. Relative Log Abundance plot was visualized to assess sample heterogeneity; samples showed a similar distribution, with medians of the boxplots close to zero and little variations, therefore sample normalization procedure was not applied.

***LC-MS analysis with HILIC***. LC-MS analyses were performed using a U3000 liquid chromatography system coupled to a Q-Exactive mass spectrometer from Thermo Fisher Scientific (Courtaboeuf, France) fitted with an electrospray source operated in the positive and negative ion modes. HPLC chromatographic separations were performed on a Sequant ZICpHILIC 5 µm, 2.1 x 150 mm (HILIC) at 15°C (Merck, Darmstadt, Germany) equipped with an on-line prefilter (Thermo Fisher Scientific, Courtaboeuf, France). Mobile phase A consisted of an aqueous buffer of 10 mM ammonium carbonate in water adjusted to pH 10.5 with ammonium hydroxide, whereas pure acetonitrile was used as solvent B. Chromatographic elutions were achieved under gradient conditions as follows: the flow rate was 200 µL/min. Elution started with an isocratic step of 2 min at 80% B, followed by a linear gradient from 80 to 40% of phase B from 2 to 12 min. The chromatographic system was then rinsed for 5 min at 0% B, and the run ended with an equilibration step of 15 min (80% B). The Exactive mass spectrometer was operated with capillary voltage at -3 kV in the negative ionization mode and a capillary temperature set at 280°C. The sheath gas pressure and the auxiliary gas pressure were set, respectively, at 60 and 10 arbitrary units with nitrogen gas. The mass resolution power of the analyzer was 50,000 m/Dm, full width at half maximum (FWHM) at m/z 200, for singly charged ions. The detection was achieved from m/z 50 to 1000.

***Processing steps and annotation.*** All raw data were manually inspected using the Qualbrowser module of Xcalibur version 2.1 (Thermo Fisher Scientific, Courtaboeuf, France). Raw files were first of all converted to mzXML format using MSConvert software. Automatic peak detection and integration were performed using the XCMS software package(Giacomoni *et al*, 2015), which returned a data matrix containing m/z and retention time values of features together with their concentrations expressed in arbitrary units (i.e., areas of chromatographic peaks). XCMS features were thereafter filtered according to the following criteria: (i) the correlation between dilution factors of QC samples and areas of chromatographic peaks (filtered variables should exhibit coefficients of correlation above 0.7 in order to account for metabolites occurring at low concentrations and which are not detected anymore in the most diluted samples), (ii) repeatability (the coefficient of variations obtained for chromatographic peak areas of QC samples should be below 30%) and (iii) ratio of chromatographic peak area of biological to blank samples above a value of 3. Optionally, if necessary, chromatographic peak areas of each variable present in the XCMS peak lists were normalized using the LOESS algorithm in order to remove analytical drift induced by clogging of the ESI source observed in the course of analytical runs.

Features were annotated by our spectral database according to accurately measured masses and chromatographic retention times obtained from ~1000 pure authentic standards (Boudah *et al*, 2014). To be identified, ions had to match at least 2 orthogonal criteria (accurately measured mass, isotopic pattern, MS/MS spectrum and retention time) to those of an authentic chemical standard analyzed under the same analytical conditions, as proposed by the Metabolomics Standards Initiative (Sumner *et al*, 2007). Features were also annotated by matching their accurate measured masses ± 10 ppm with theoretical masses contained in biochemical and metabolomic databases by using an informatics tool developed in R language. The databases used were the Kyoto Encyclopedia of Genes and Genomes (KEGG) (Kanehisa, 2000), the Human Metabolome Database (HMDB) (Wishart *et al*, 2022) and METLIN (Smith *et al*, 2005).

**MS2. ML analyses on plasma samples metabolomics data**

Metabolomics data for 81 plasma samples were used as input to various classifiers. Data was normalized separately for the C18 and HILIC datasets by dividing all raw signal values by the average values found for the mild samples (*cf.* 'Covid_Data_MS_ML' Supplementary file). The two normalized datasets were merged and fed to the ML classifiers. The merged dataset comprised 624 MS signals (423 from C18 and 201 from HILIC). Additionally, we reduced the merged dataset selecting only MS signals corresponding to metabolites crossing the *E. coli* membrane. These medium metabolites were those found in the exchange reaction of iML1515 model. The mapping between the MS metabolites/ions and iML1515 exchange metabolites was carried out using their InChI code which was found in the HMDB database from their names. The reduced medium dataset comprised 56 metabolites.

As classifiers, we used the SVM, MLP and XGB classifiers from the SciKitLearn (Pedregosa *et al*, 2012) library. Default parameters were used in all cases and for MLP, the neural network comprised one hidden layer with a size twice the input size. MLP was trained for 10000 epochs with early stopping and the Adam optimizer. For all classifiers 20-fold cross validation was performed. Standard deviations were acquired iterating the cross-validation process 5 times.

Results are presented in Supplementary Figure **S1**, these were obtained running the above classifiers after feature selection (Supplementary Methods **MS3**).

**MS3. Feature selection for plasma samples classification**

When performing COVID-19 plasma samples classification, accuracies were calculated after feature selection. To find the final set of features (*i.e.* the set of selected MS signals or the set of selected OD time points) we removed features one at a time, run the classification, and at each iteration the deleted feature is the one leading to the highest accuracy after deletion. The accuracies following selected features for COVID-19 mild vs. severe classification using MS data are shown in Supplementary Figure **S7** (and Supplementary file 'Covid_Data_MS_ML'). The accuracies following selected features for COVID-19 negative vs. positive and mild vs. severe classification with *E. coli* physical reservoir predictions data are shown in Supplementary Figure **S8** (and Supplementary file 'Reservoir_Covid).

### Supplementary Tables

**Table S1. *E. coli* reservoir COVID-19 classification using lag time, growth rate and $OD_{MAX}$.**
*XGB classification using as input lag time, growth rate, and $OD_{MAX}$. Lag time, growth rate, and $OD_{MAX}$ are calculated following as detail in Materials and Methods sections **M6.3.** Results are reported for 20-fold cross validation aggregated sets.*

| Feature | Negative vs. Positive<br>Mean accuracy ± dev (for 3 repeat) | Mild vs. Severe<br>Mean accuracy ± dev (for 3 repeat) |
|---|---|---|
| Growth Rate | 0.72±0.005 | 0.58±0.011 |
| Lag Time<br>Growth Rate<br>$OD_{MAX}$ | 0.85±0.025 | 0.65±0.006 |

**Table S2. *E. coli* reservoir COVID-19 classification for several classifiers and cross-validation folds.**
*Classification is performed taking as input growth curves, i.e., OD values after selecting time points among 48 (from 0 to 8 hours, cf. Supplementary Methods **MS3**). The results shown for XGB and 20 folds are those presented in **Fig. 2** (main manuscript).*

| Method | Cross validation fold | Negative vs. Positive<br>Mean accuracy ± dev (for 3 repeat) | Mild vs. Severe<br>Mean accuracy ± dev (for 3 repeat) |
|---|---|---|---|
| SVM | 5 | 0.84±0.004 | 0.68±0.030 |
| | 20 | 0.85±0.001 | 0.73±0.020 |
| | LOO | 0.86±0.001 | 0.70±0.001 |
| MLP | 5 | 0.90±0.029 | 0.77±0.030 |
| | 20 | 0.92±0.016 | 0.79±0.010 |
| | LOO | 0.93±0.008 | 0.80±0.015 |
| XGB | 5 | 0.94±0.005 | 0.77±0.017 |
| | 20 | 0.95±0.005 | 0.86±0.001 |
| | LOO | 0.95±0.005 | 0.85±0.006 |

**Table S3. *E. coli* reservoir growth dynamics statistical analysis.**

*This table presents the statistical analysis comparing E. coli growth dynamics between negative vs. positive and mild vs. severe conditions using Generalized Additive Mixed Models (GAMMs)* (Wood, 2017). *The **t-value** is calculated as the ratio of the estimator to its standard error. It is used to assess statistical significance based on the Student's t-distribution. The t-value indicates the size of the effect relative to its variability. **EDF** (Effective Degrees of Freedom) reflects the complexity of the spline smooth term in the model. A higher EDF indicates a more flexible, or "wigglier," curve. For example, an EDF of 8.8 suggests the best fit is close to a 9th-order polynomial, though penalized to reduce overfitting and ensure smoothness. **Ref.df** (Reference Degrees of Freedom) is analogous to the degrees of freedom in a traditional parametric model, serving as a baseline for statistical comparison. The **F-statistic** measures the effect of a term relative to its variance, based on the Fisher distribution. Like the t-value, it is used to assess significance but is typically applied in the context of ANOVA. The **t-value** and **F-statistic** are outputs of the model and are used to compute **p-values**, which indicate the statistical significance of each term in the model. **s(Time)** stands for spline(Time) - smooth spline function for Time variable.*

<table>
<tr><td rowspan="8">Group Positive vs Group Negative (R2 adjusted = 0.791)</td><td colspan="5">Parametric coefficients</td></tr>
<tr><td></td><td>Estimate</td><td>Std. Error</td><td>t value</td><td>Prob(>|t|)</td></tr>
<tr><td>Intercept</td><td>0.088991</td><td>0.004312</td><td>20.640</td><td><2e-16*</td></tr>
<tr><td>Group Positive</td><td>-0.020407</td><td>0.005046</td><td>-4.044</td><td>5.37e-05*</td></tr>
<tr><td colspan="5">Approximate significance of smooth terms</td></tr>
<tr><td></td><td>edf</td><td>Ref.df</td><td>F</td><td>p-value</td></tr>
<tr><td>s(Time):Group Negative</td><td>8.856</td><td>8.856</td><td>214.0</td><td><2e-16*</td></tr>
<tr><td>s(Time):Group Positive</td><td>8.915</td><td>8.915</td><td>425.6</td><td><2e-16*</td></tr>
<tr><td rowspan="8">Group Mild vs Group Severse (R2 adjusted = 0.783)</td><td colspan="5">Parametric coefficients</td></tr>
<tr><td></td><td>Estimate</td><td>Std. Error</td><td>t value</td><td>Prob(>|t|)</td></tr>
<tr><td>Intercept</td><td>0.067837</td><td>0.002371</td><td>28.606</td><td><2e-16*</td></tr>
<tr><td>Group Severe</td><td>0.001516</td><td>0.003377</td><td>0.449</td><td>0.654</td></tr>
<tr><td colspan="5">Approximate significance of smooth terms</td></tr>
<tr><td></td><td>edf</td><td>Ref.df</td><td>F</td><td>p-value</td></tr>
<tr><td>s(Time):Group Mild</td><td>8.818</td><td>8.818</td><td>227.7</td><td><2e-16*</td></tr>
<tr><td>s(Time):Group Severe</td><td>8.886</td><td>8.886</td><td>275.9</td><td><2e-16*</td></tr>
</table>

*Significant values

**Table S4. Learners architectures and hyperparameters.**
*Most hyperparameters values are the default values for the functions found in ScikitLearn, XGboost or Tensorflow. Baseline comparators are not integrated in RC frameworks but used for performance comparisons in* ***Figs. 4-6****.*

| Role (dataset) | Method | Library Function | Hyperparameters | Epochs k-fold CV |
|---|---|---|---|---|
| features-to-medium mapper (ML datasets) | MLP regression | Tensorflow Dense | Hidden layer sizes = (280, 280, 280), Adam optimizer, Learning rate: $10^{-3}$, Batch Size: 16, Activation: 'Gumbel-Softmax' | 1000<br>5 |
| AMN (100-medium set) | MLP regression | Tensorflow Dense | Hidden layer size = 500, Adam optimizer, Learning rate: $10^{-3}$, Batch Size: 10, Activation: 'ReLU' | 1000<br>10 |
| Readout (COVID datasets) | SVM classification | ScikitLearn SVC | Kernel: 'polynomial' | 10000<br>20 |
| Readout (COVID datasets) | MLP classification | Tensorflow Dense | Hidden layer size = Input size / 2, Adam optimizer, Learning rate: $10^{-3}$, Batch Size: 5, Activation: 'sigmoid' | 1000<br>20 |
| Readout (COVID datasets) | XGB classification | XGBoost XGBClassifier | n_estimators: 100, learning_rate: 0.1 | n/a<br>20 |
| Readout (ML datasets) | Linear regressor | ScikitLearn linear_model Ridge | alpha: 1.0, solver = 'auto', tol: 0.0001 | n/a<br>5 |
| Readout (ML datasets) | Linear classifier | ScikitLearn linear_model RidgeClassifier | alpha: 1.0, solver = 'auto', tol: 0.0001, class_weigth: None | n/a<br>5 |
| Baseline comparator (ML datasets) | MLR | ScikitLearn LinearRegression | n/a | n/a<br>5 |
| Baseline comparator (ML datasets) | SVM classification | ScikitLearn SVC | Kernel: 'linear' | 10000<br>5 |
| Baseline comparator (ML datasets) | MLP regression | ScikitLearn MLPRegressor | Adaptive Learning rate, Early stopping, Adam optimizer, Activation: 'ReLU' | 10000<br>5 |
| Baseline comparator (ML datasets) | MLP classification | ScikitLearn MLPClassifier | Adaptive learning rate, Early stopping, Adam optimizer, Activation: 'sigmoid' | 10000<br>5 |
| Baseline comparator (ML datasets) | XGB regression | XGBoost XGBRegressor | n_estimators: 1, learning_rate: 0.3 | n/a<br>5 |
| Baseline comparator (ML datasets) | XGB classification | XGBoost XGBClassifier | n_estimators: 100, learning_rate: 0.1 | n/a<br>5 |

**Table S5. Genome Scale Metabolic Models (GEMs).**
*The table provides the list of GEMs used in the study with their main characteristics. Mono-directional reaction are required for AMN implementation. The number of mono directional reactions is calculated as in* (Faure *et al*, 2023).

| BiGG ID | Short name | Organisms | Meta-bolites | Reac tions | Mono directional reaction | Genes | Reference |
|---|---|---|---|---|---|---|---|
| iEK1008 | *M. tuberculosis* | *Mycobacterium tuberculosis* H37Rv | 998 | 1226 | 1597 | 1008 | Kavvas *et al.* *DOI: 10.1186/s12918-018-0557-y* |
| iIT341 | *H. pylori* | *Helicobacter pylori* 26695 | 485 | 554 | 793 | 339 | Thiele *et al.* *DOI: 10.1128/JB.187.16.5818-5830.2005* |
| iJN1463 | *P. putida* | *Pseudomonas putida* KT2440 | 2153 | 2927 | 4034 | 1462 | Nogales *et al.* *DOI: 10.1111/1462-2920.14843* |
| iML1515 | *E.coli* | *E. coli* str. K-12 substr. MG1655 | 1877 | 2712 | 3682 | 1516 | Monk *et al.* *DOI: 10.1038/nbt.3956* |
| iMM904 | *S. cerevisiae* | *Saccharomyces cerevisiae* S288C | 1226 | 1577 | 2226 | 905 | Mo *et al.* *DOI: 10.1186/1752-0509-3-37* |
| iPC815 | *Y.pestis* | *Yersinia pestis* CO92 | 1552 | 1961 | 2767 | 815 | Charusanti *et al.* *DOI: 10.1186/1752-0509-5-163* |
| iYO844 | *B. subtilis* | *Bacillus subtilis subsp. subtilis* str. 168 | 990 | 1250 | 1806 | 844 | Oh *et al.* *DOI: 10.1074/jbc.M703759200* |
| iYS1720 | *Salmonella* | *Salmonella* pan-reactome | 2436 | 3357 | 4451 | 1707 | Seif *et al.* *DOI: 10.1038/s41467-018-06112-5* |
| iYS854 | *S. aureus* | *Staphylococcus aureus subsp. aureus* USA300_TCH1516 | 1335 | 1455 | 2086 | 866 | Seif *et al.* *DOI: 10.1371/journal.pcbi.1006644* |
| iCN718 | *Acinetobacter baumannii* | *Acinetobacter baumannii* AYE | 888 | 1015 | 1438 | 709 | Norsigian *et al.* *DOI: 10.3389/fgene.2018.00121.* |
| iNF517 | *Lactococcus lactis* | *Lactococcus lactis* subsp. cremoris MG1363 | 650 | 754 | 1079 | 516 | Flahaut *et al.* *DOI: 10.1007/s00253-013-5140-2* |

**Table S6. Regression problems extracted from OpenML.**
*List of all regression problems used in the study.*

| Problem | Instances | Features | Reference |
|---|---|---|---|
| white wine | 4898 | 11 | www.openml.org/search?type=data&sort=runs&id=40498&status=active |
| kin8nm | 8192 | 8 | www.openml.org/search?type=data&sort=runs&id=189&status=active |
| cars | 392 | 7 | www.openml.org/search?type=data&sort=runs&id=21&status=active. |
| airfoil noise | 1503 | 5 | www.openml.org/search?type=data&sort=runs&id=44957&status=active |
| fish toxicity | 908 | 6 | www.openml.org/search?type=data&sort=runs&id=44970&status=active |
| space ga | 3107 | 6 | www.openml.org/search?type=data&sort=runs&id=507&status=active |
| concrete | 1030 | 8 | www.openml.org/search?type=data&sort=runs&id=44959&status=active |
| grid stability | 10000 | 12 | www.openml.org/search?type=data&sort=version&status=any&order=asc&id=43007 |
| cpu activity | 8192 | 21 | www.openml.org/search?type=data&sort=runs&id=197&status=active |
| energy efficiency | 768 | 8 | www.openml.org/search?type=data&sort=runs&id=1472&status=active |

#### Supplementary Figures

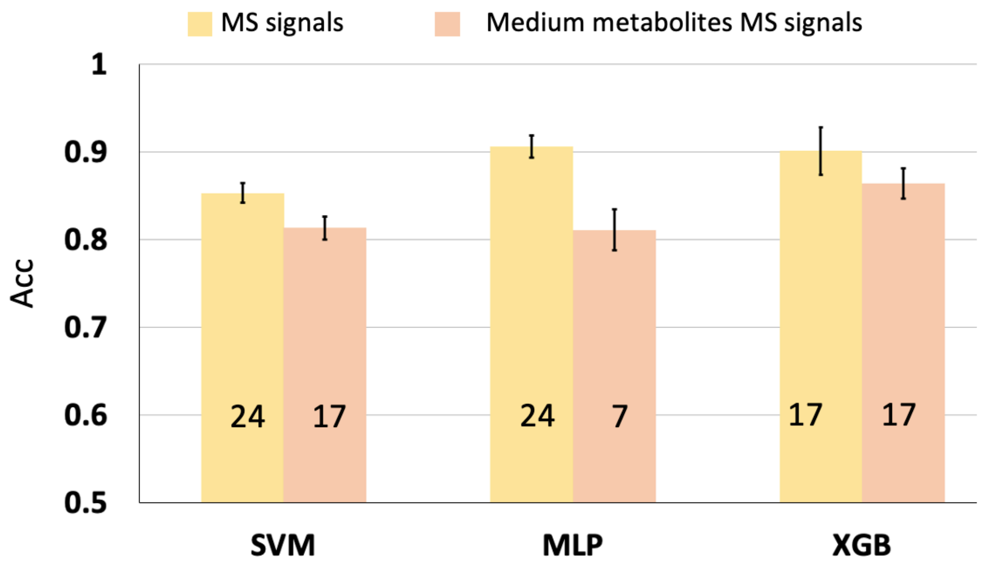

**Figure S1. Classification of COVID-19 plasma samples with metabolomics data.**
*(a) Classifying (mild vs. severe) COVID-19 with metabolomics data. MS signals: accuracies using all 624 metabolite signals detected by MS for 81 patients. Medium metabolites MS signals: accuracies using 56 MS signals for metabolites expected to cross the E. coli cell wall. Accuracies are calculated for 20-fold cross validation and using feature selection (cf. section* ***MS3*** *and Supplementary Figure* ***S7****). The numbers indicated on the bar charts are the numbers of features selected. The parameters used for the classifiers (SVM, XGB and MLP) are given in Supplementary Table* ***S4****. All data are in Supplementary file 'Covid_Data_MS_ML'.*

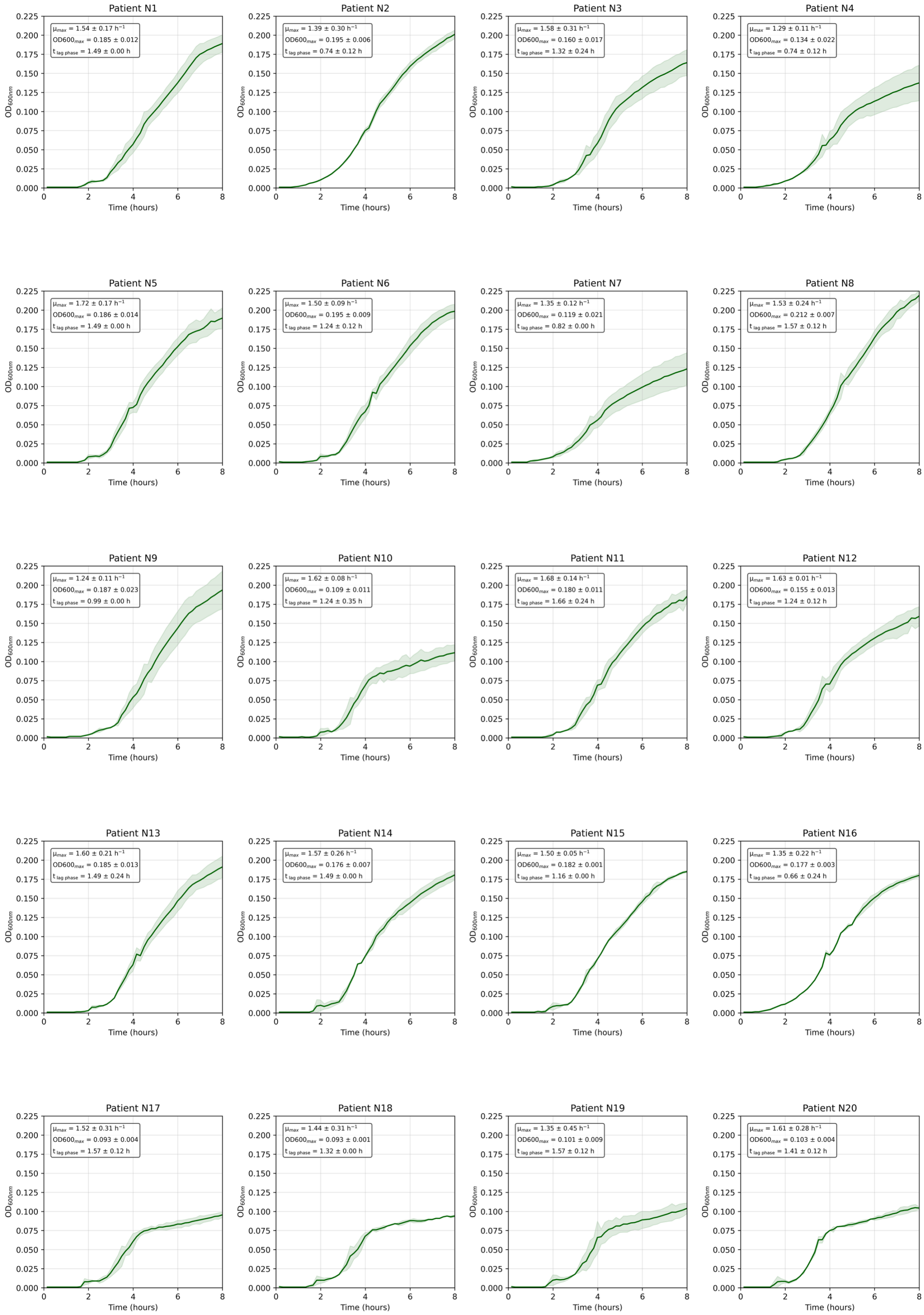

**Figure S2. *E. coli* growth curves cultured in COVID-19 negative plasma samples.**
*Data acquired in technical duplicates. Data are in the Supplementary file 'Reservoir-Covid'. The captions indicate the maximal growth rate, the $OD600_{max}$ (mean of the 5 highest optical density measurements taken at 600 nm) and the lag phase duration for each sample.*

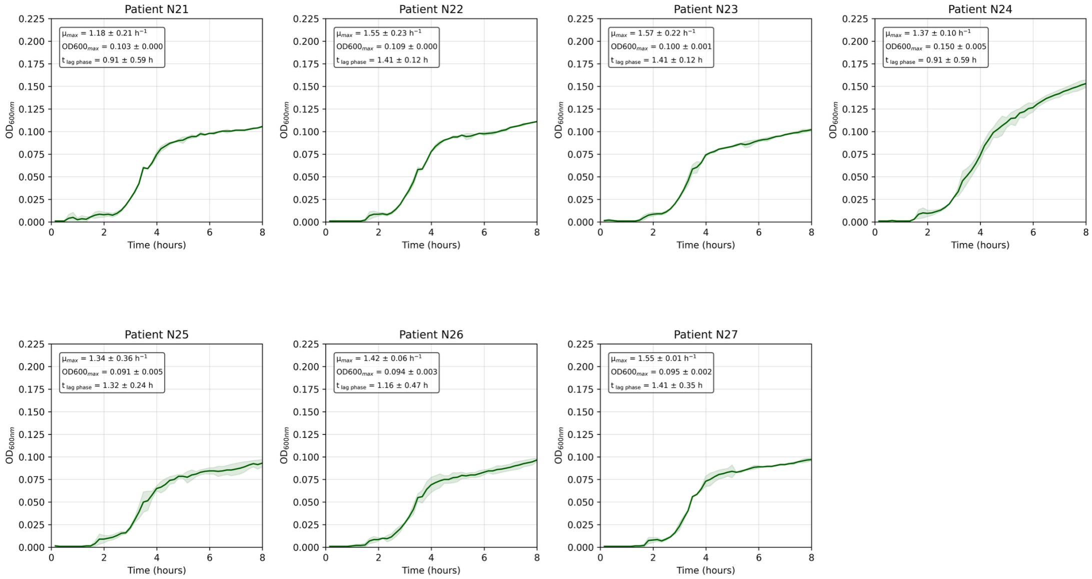

**Figure S2 (cont). *E. coli* growth curves cultured in COVID-19 negative plasma samples.** *Data acquired in technical duplicates. Data are in the Supplementary file 'Reservoir-Covid'. The captions indicate the maximal growth rate, the $OD600_{max}$ (mean of the 5 highest optical density measurements taken at 600 nm) and the lag phase duration for each sample.*

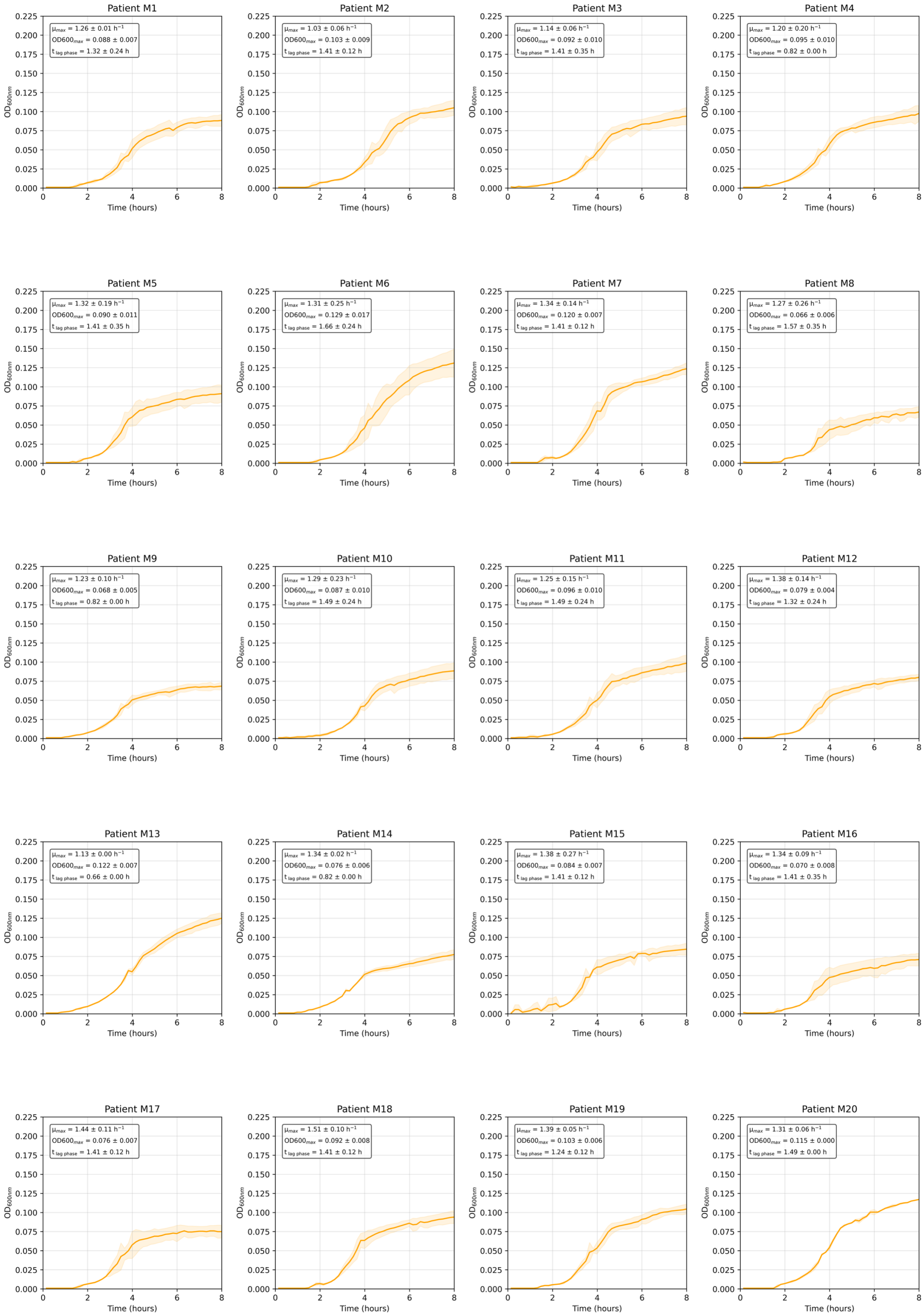

**Figure S3. *E. coli* growth curves cultured in COVID-19 plasma samples from the mild group.**

*Patients were sampled prior to developing a mild form of the disease. Data acquired in technical duplicates. Data are in the Supplementary file 'Reservoir-Covid'. The captions indicate the maximal growth rate, the OD600$_{max}$ (mean of the 5 highest optical density measurements taken at 600 nm) and the lag phase duration for each sample.*

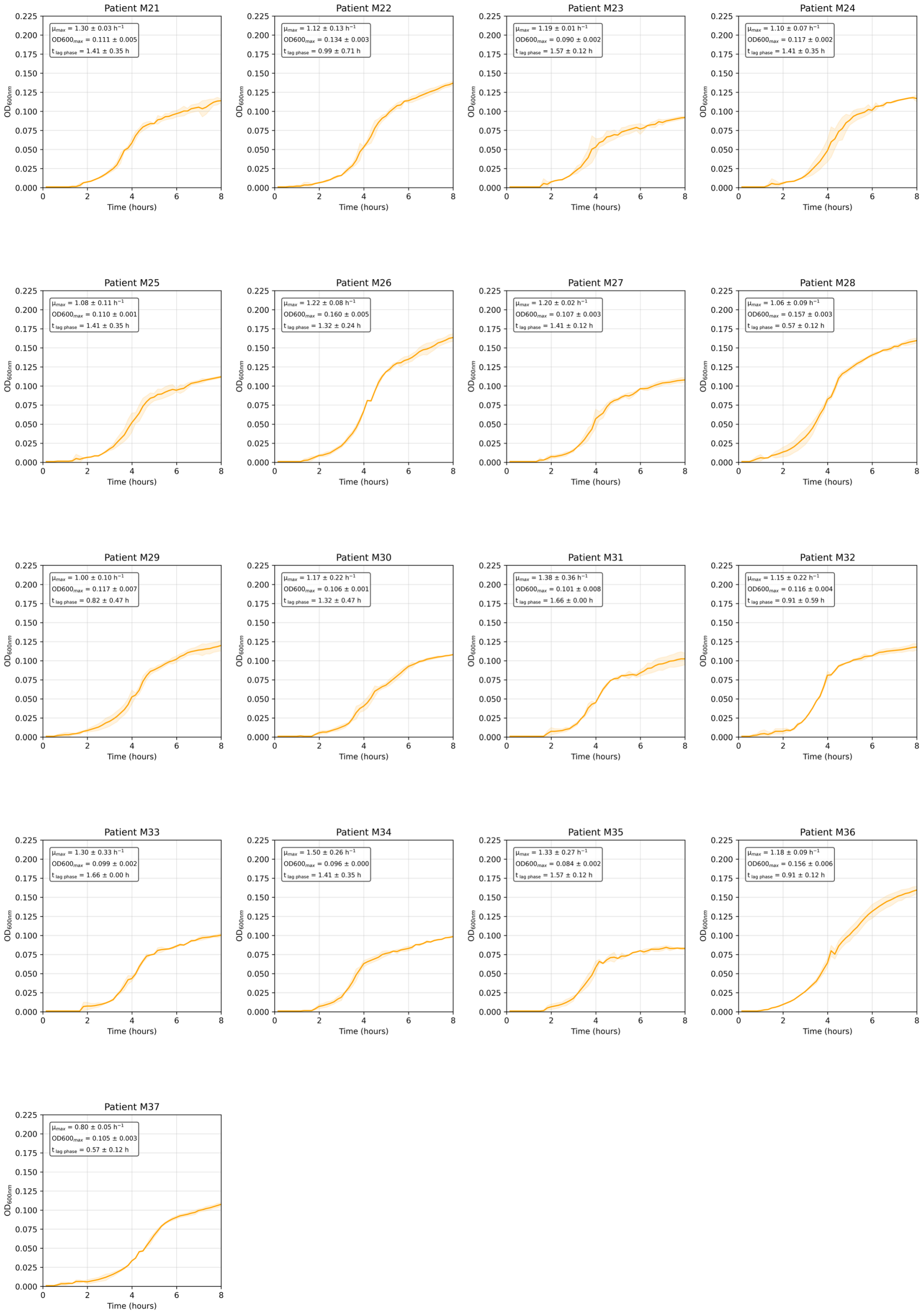

**Figure S3 (cont).** ***E. coli*** **growth curves cultured in COVID-19 plasma samples from the mild group.**

*Patients were sampled prior to developing a mild form of the disease. Data acquired in technical duplicates. Data are in the Supplementary file 'Reservoir-Covid'. The captions indicate the maximal growth rate, the $OD600_{max}$ (mean of the 5 highest optical density measurements taken at 600 nm) and the lag phase duration for each sample.*

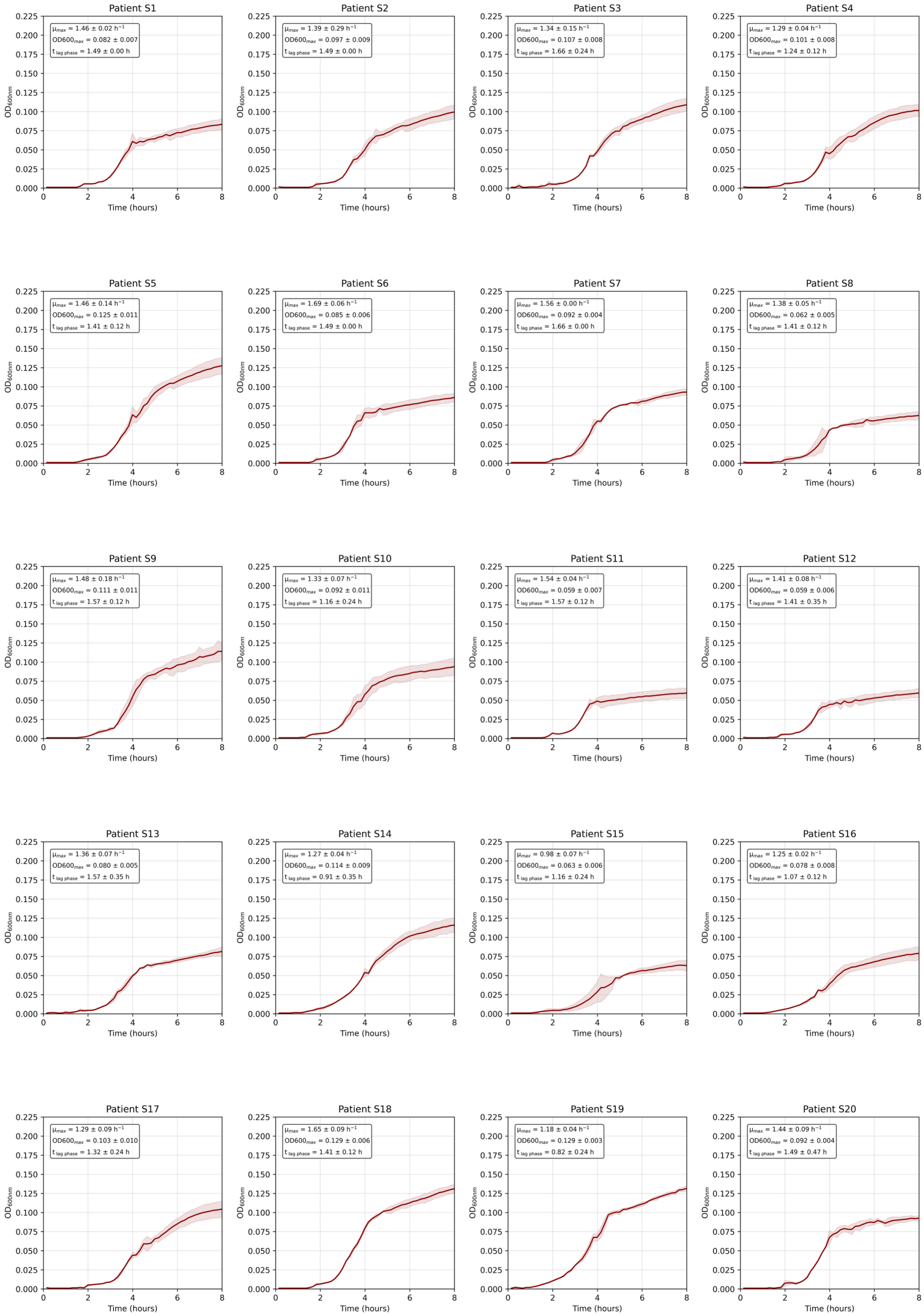

**Figure S4. *E. coli* growth curves cultured in COVID-19 plasma samples from the severe group.**

*Patients were sampled prior to developing a severe form of the disease. Data acquired in technical duplicates. Data are in the Supplementary file 'Reservoir-Covid'. The captions indicate the maximal growth rate, the OD600$_{max}$ (mean of the 5 highest optical density measurements taken at 600 nm) and the lag phase duration for each sample.*

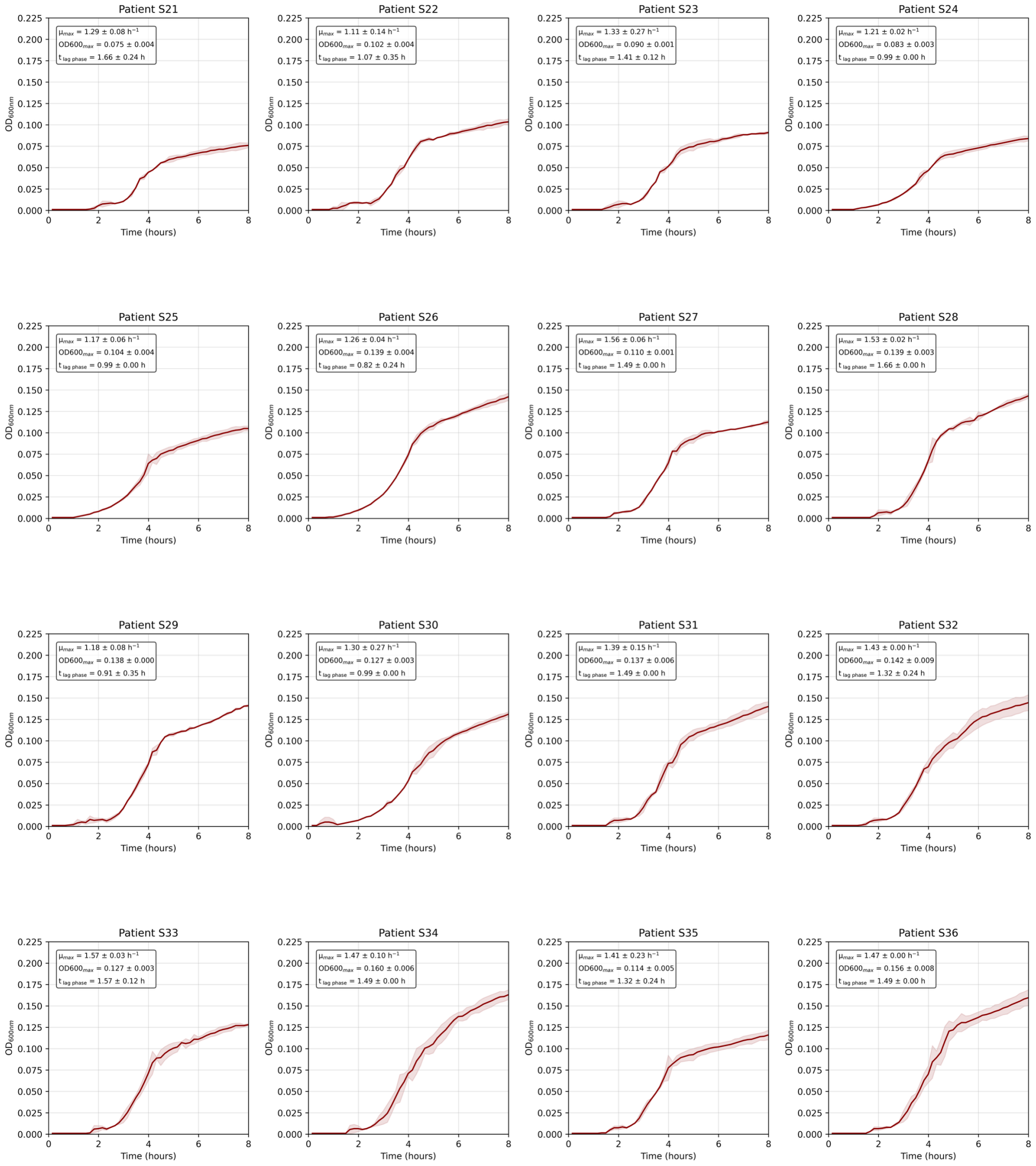

**Figure S4 (cont). *E. coli* growth curves cultured in COVID-19 plasma samples from the severe group.**

*Patients were sampled prior to developing a severe form of the disease. Data acquired in technical duplicates. Data are in the Supplementary file 'Reservoir-Covid'. The captions indicate the maximal growth rate, the $OD600_{max}$ (mean of the 5 highest optical density measurements taken at 600 nm) and the lag phase duration for each sample.*

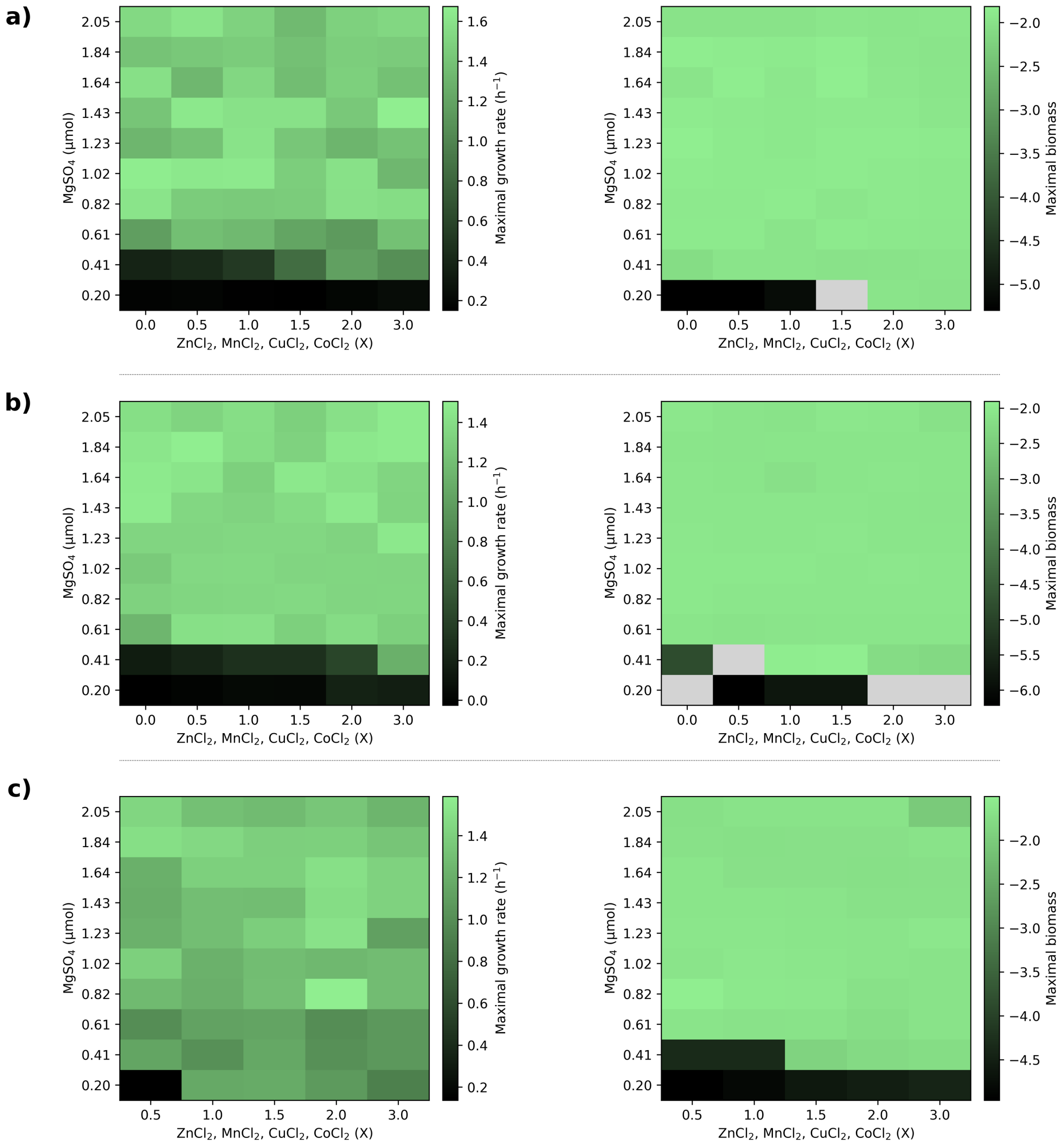

**Figure S5. Growth optimization of *E. coli* in plasma-EDTA samples.**
*Escherichia coli K-12 MG1655 was cultured in plasma-EDTA samples from COVID-19-negative patients, supplemented with increasing quantities of $MgSO_4$, $ZnCl_2$, $MnCl_2$, $CuCl_2$, and $CoCl_2$. y-axis values represent the quantity of $MgSO_4$ in µmol supplemented in each condition. x-axis values represent multiples of "X", a fixed mixture used as a reference standard, composed of 3.08 nmol $ZnCl_2$, 0.38 nmol $MnCl_2$, 0.21 nmol $CuCl_2$, and 0.04 nmol $CoCl_2$. $MgSO_4$ was varied independently, as magnesium is the only ion among those tested that is required for E. coli growth. Heatmaps display the maximal growth rate (left panels) and maximal biomass (right panels; natural logarithm of $OD_{600}$ measured at the onset of stationary phase) for each condition. Grey boxes indicate conditions where growth was not observed and maximal biomass could not be determined. Panels a–c represent biological replicates 1, 2, and 3, respectively.*

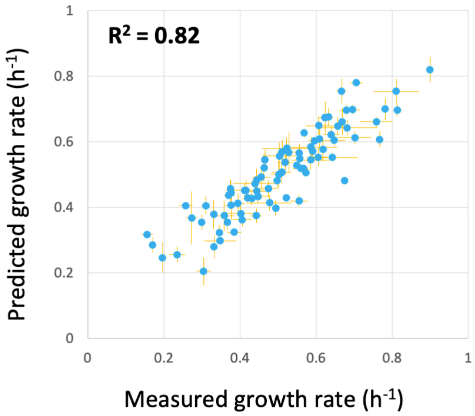

**Figure S6. AMN training results.**
*Correlation between AMN predicted and measured growth rates. AMN was trained on 100 media with measured growth rates as in* (Faure *et al*, 2023). *All points were acquired on cross-validation sets (10-fold); error bars (both measured and predicted) represent standard deviations over 3 and 50 repeats respectively. Data are in the Supplementary file 'Ecoli_Data'*

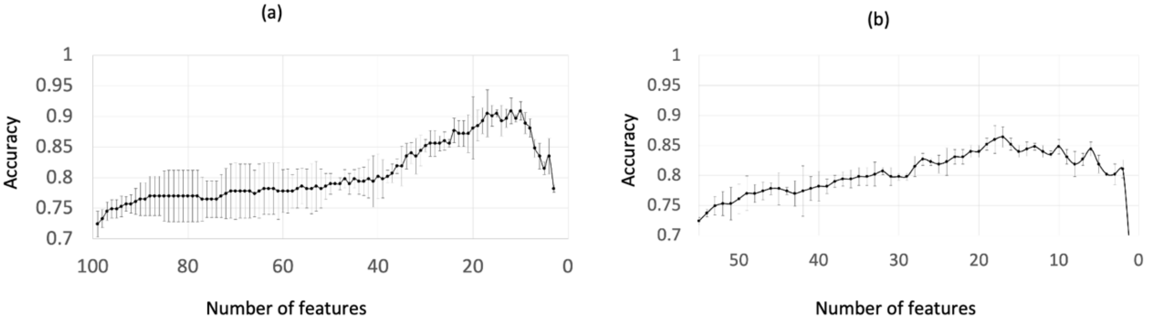

**Figure S7. Features selection for COVID-19 samples classification with MS data.**
*(a) For all MS signals. The curve shows the accuracies using XGB classifier for the 100 most important features (among 620 MS signals) removing at each step the feature leading to the highest accuracy after deletion . (b) Same as in (a) but for all 52 MS signals corresponding to medium metabolites. Data are in the Supplementary file 'Covid_Data_MS_ML.*

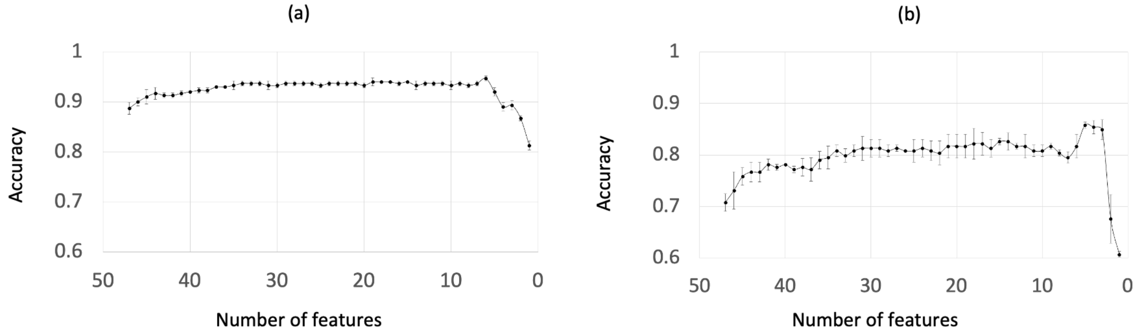

**Figure S8. Features selection for COVID-19 samples classification with *E. coli* reservoir.**
*The curve shows the accuracies using XGB classifier removing at each step the feature (time point) leading to the highest accuracy after deletion (a) For negative vs. positive classification. (b) For mild vs. severe classification. Data are in the Supplementary file 'Reservoir-Covid'.*

### Other Supporting Materials

Data_S1_Covid_Data_MS_ML: patient data for BIOMARCOVID cohort, MS data on cohort plasma samples, mild vs. severe cross-validation results using various classifiers

Data_S2_Reservoir_Covid: patient data for BIOMARCOVID cohort, optical density data growing E. coli MG1655 on negative, mild, and severe BIOMARCOVID plasma samples.

Data_S3_Ecoli_Data: Training results for AMN trained with growth rates on 100 medium library, AMN prediction for 280 media and *E. coli* strain MG1655 growth curves for 280 media

Data_S4_Reservoir_Classification : Cross validation results for 8 classification tasks with *E. coli* reservoir.

Data_S5_Reservoir_Regression: Cross validation results for 10 regression tasks with *E. coli* reservoir and for 10 *in-silico* GEM reservoirs.
